# A collicular somatostatinergic circuit gates sensory access to action

**DOI:** 10.64898/2026.09.14.751531

**Authors:** Daniel de Malmazet, Letizia Mariotti, Yuanxin Zhang, Daniel Welch, Lynn Geyer, Antonio Falasconi, Fabio Morgese, Marco Tripodi

## Abstract

Animals are exposed to far more sensory information than can be converted into action and must prioritise behaviourally salient events. Although salience is commonly studied as modulation of sensory representations, stimulus prioritisation may also arise at premotor stages, through mechanisms that regulate the readiness of action-generating circuits to be recruited by sensory input. The superior colliculus links visual signals to spatially directed orienting and contains Pitx2-expressing spatial-motor modules that coordinate spatially targeted actions, providing a substrate for premotor regulation of sensory access to action. However, the circuit elements that regulate this access remain unknown. Here we identify an intersectionally targeted population of somatostatin-expressing inhibitory neurons in the mouse superior colliculus that is, at the population level, suppressed during orienting movements and visual stimulation. These neurons provide input to Pitx2-expressing spatial-motor modules, and their optogenetic silencing increased retinally evoked firing in Pitx2 neurons, showing that somatostatinergic inhibition constrains the visual recruitment of collicular motor output. Increasing somatostatinergic tone reduced interception of low- and intermediate-contrast visual targets while sparing responses to high-contrast targets, consistent with graded control of sensory access to action. Cholecystokinin-expressing inhibitory neurons and multiple cortical and subcortical regions provide anatomical input to the somatostatinergic population, identifying candidate routes for local and context-dependent regulation. Together, these findings establish somatostatinergic inhibition as a premotor control point that regulates the readiness of spatial-motor circuits for sensory recruitment and identify a circuit architecture through which the behavioural impact of salient sensory events could be regulated downstream of sensory encoding.

## Introduction

Animals must transform only a subset of environmental events into action^1^. The behavioural impact of salient events is often studied through changes in sensory representations^2,3^, but an additional control point may lie downstream, at the interface between sensory input and motor output. The same sensory signal could therefore have different behavioural consequences depending on the readiness of premotor circuits to be recruited. A circuit that regulates the excitability of action-generating neurons could alter the behavioural impact of salience without requiring the sensory representation itself to be amplified^4^.

The superior colliculus (SC) has been shown to carry information concerning the onset of biologically relevant events^5–8^. Across vertebrates, it integrates sensory signals with commands for spatially directed movements^9–16^. Studies of natural visuomotor behaviour further indicate that the recruitment of collicular neuronal assemblies can link perception to the initiation of action^11,16–25^. In mice, Pitx2-expressing projection neurons form spatial-motor modules that control orienting^9,15^. Dynamic inhibition of these output neurons could therefore regulate their readiness to be recruited by sensory input, providing a premotor mechanism through which stimulus salience could influence action. Although intrinsic inhibition in the motor SC is well established^18,26,27^, the molecularly accessible inhibitory populations that implement this control and their relationship to motor output modules remain unresolved^28,29^. The relative lack of knowledge about the role of intrinsic collicular inhibitory neurons contrasts with accumulating evidence of molecularly^30–33^ and functionally^34–38^ distinct types of inhibitory neurons in the cortex, supporting several computational functions including gating and gain modulation^39,40^.

Here, we combined freely moving and head-fixed calcium imaging, intersectional genetics, circuit tracing, slice electrophysiology and behavioural manipulation to identify inhibitory populations associated with orienting and to test how one of these populations controls the recruitment of Pitx2 spatial-motor modules by retinal input.

We find that the intersectionally targeted Sst inhibitory neurons are negatively coupled to movement and visual stimulation, provide input to Pitx2 neurons, and silencing the population increases retinally evoked Pitx2 firing. Conversely, increasing Sst inhibitory activity preferentially reduces orienting to lower-contrast visual targets. Anatomical mapping further identifies local Cck inhibitory neurons and several extra-collicular regions as candidate regulators of this inhibitory node. These findings establish somatostatinergic inhibition as a premotor control point for sensory access to action and identify the motor readiness of spatial-motor modules as a mechanism through which the behavioural impact of salient sensory events can be regulated.

## Results

### Freely moving imaging reveals opposing inhibitory dynamics during orienting behaviour

To examine inhibitory activity during natural orienting, we performed single-photon *in vivo* calcium imaging in freely moving mice using a head-mounted miniendoscope together with Head-in-Body kinematic tracking^41^ (Fig. 1A). The implant did not measurably alter the range or structure of head movements (Fig. 1B and Supplementary Fig. 1). In wild-type mice, approximately 20% of recorded motor-SC neurons were tuned to particular head displacements, predominantly contralateral rotations, and most retained their tuning in darkness (Supplementary Fig. 2).

**Fig. 1:**
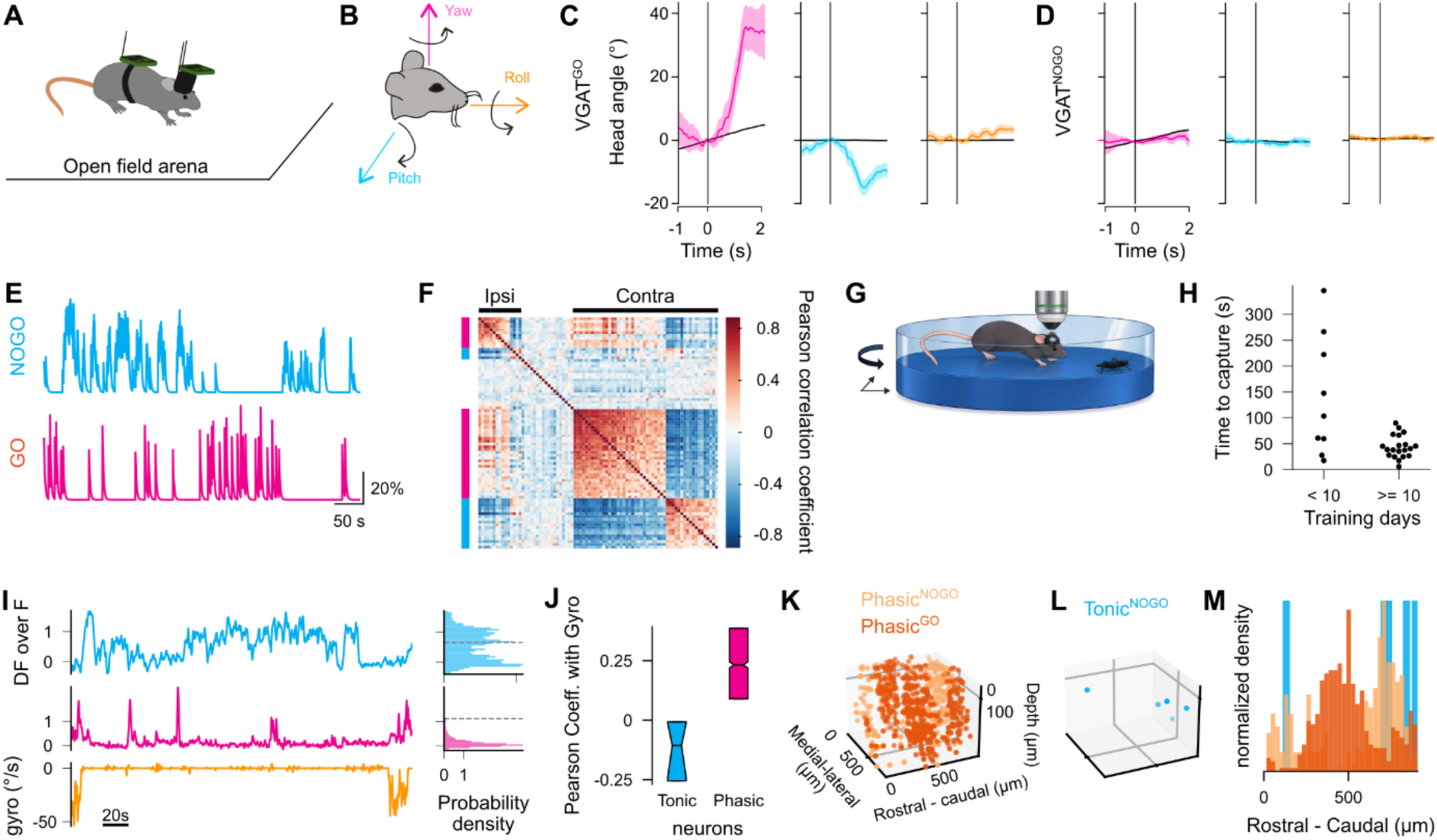
Collicular inhibitory activity contains movement-coupled and movement-suppressed components. **(A)** Mice moved freely inside an open field arena, carrying a head-mounted miniendoscope coupled with head and body inertial sensors. **(B)** Schematic representing the three axes of head rotations, roll (orange), yaw (magenta) and pitch (cyan). **(C)** Example head tuning curves of an inhibitory neuron that showed a tuning for a particular head rotation. We named this type of cells VGAT^GO^. Traces are the spike triggered average of head displacement along yaw (magenta), pitch (cyan), roll (orange). Spikes were inferred from the deconvolved calcium traces. Shaded areas represent the standard error of the mean. Black curves correspond to the spike triggered average obtained after shuffling the spike times. The vertical lines at time 0 correspond to the onset of calcium events. **(D)** Same as C for an inhibitory cell that did not show tuning for a specific head rotation. Since this cell fired when the mouse did not turn its head, we named this type of cells VGAT^NOGO^. **(E)** Example calcium traces of a VGAT^GO^ (in magenta) and VGAT^NOGO^ (in cyan) cells showing the anticorrelated relationship that we observed between the two types of inhibitory neurons. Y axis corresponds to the percentage of variation in the calcium fluorescent signal. (**F)** Correlation matrix showing the Pearson correlation coefficients for pairs of inhibitory neurons. VGAT^GO^ and VGAT^NOGO^ neurons are indicated by the magenta and cyan lines on the left, respectively. Black bars on top indicate the neurons tuned to ipsi- and contra-lateral movements (N=114 neurons). **(G)** Experimental set up for *in vivo* two-photon calcium imaging of collicular neurons. Mice were head-fixed on a floating platform. Mice were trained to catch a cricket while on the platform. **(H)** Time in seconds it took mice to catch the cricket following its placement on the platform. Each black dot corresponds to a behaviour session. Mean ± standard deviation, before 10 days of training: 139.09 ± 108.86, after 10 days: 44.17 ± 21.26. N_mice_ = 6, N_sessions_ = 29. **(I)** Example calcium traces recorded with *in vivo* two-photon calcium imaging of a tonic and a phasic neuron in cyan and magenta, respectively. The synchronised platform rotation speed is at the bottom in orange. Histograms on the right correspond to the distribution of calcium signal variations (DF over F) across time. Y axes are the same for the traces and the corresponding histograms. Dotted lines mark the midrange of the DF over F distributions. **(J)** Pearson correlation coefficients between the platform rotation speed and the calcium traces of tonic and phasic neurons. Boxplots show median and [interquartile range]; Tonic neurons: −0.11 [−0.25, −0.01]; Phasic neurons: 0.23 [0.09, 0.39]. Notches indicate the 95% confidence interval of the median bootstrapped 10000 times. **(K)** Anatomical positions of Phasic^GO^ and Phasic^NOGO^ neurons from a volumetric recording of the SC. **(L)** Anatomical positions of Tonic^NOGO^ neurons from the same volumetric recording of the SC as in K. **(M)** Density of Tonic^NOGO^ (blue), Phasic^GO^ (red) and Phasic^NOGO^ (beige) neurons along the rostro-caudal axis of the imaged SC.

We next compared genetically targeted glutamatergic and GABAergic populations by expressing GCaMP6s in VGLUT2-Cre or VGAT-Cre mice (6 mice, 578 neurons). Motor-tuned VGLUT2^ON^ neurons resembled the movement-related units described previously. Within the VGAT^ON^ population, some neurons were active around specific head rotations, whereas others were preferentially active during periods of little or no head movement (Fig. 1C-F). We refer to these descriptive activity patterns as VGAT^GO^ and VGAT^NOGO^, respectively.

Thus, inhibitory SC activity contains movement-coupled and movement-suppressed patterns that are associated with opposing behavioural states. These observations motivated a higher-throughput analysis of inhibitory population dynamics.

### Population imaging resolves distinct movement-related inhibitory activity patterns

To sample larger inhibitory populations across multiple fields of view, we performed two-photon *in vivo* calcium imaging through a cranial window while head-fixed mice moved an air-supported circular platform. Platform rotation reoriented the animal’s body relative to the head and was measured continuously with an inertial sensor (Fig. 1G).

As a behavioural validation of the platform, mice learned to orient towards, approach and capture a live cricket while head-fixed, reaching capture latencies comparable to those reported in freely moving mice after training (Fig. 1H and Supplementary Video 1)^42–45^. This experiment established that the platform supported coordinated visually guided orienting but was not used to classify neuronal responses.

We then recorded VGAT^ON^ neurons during spontaneous platform rotations in complete darkness, thereby minimizing visually evoked activity (Supplementary Fig. 3A-B). Across 13 recordings from two mice, most neurons changed activity around movement, although the temporal profiles were heterogeneous. A small subset showed sustained activity during immobility and was negatively coupled to rotation, whereas the majority displayed phasic activity associated with movement (Fig. 1I-J and Supplementary Fig. 3C). We refer to these profiles as tonic and phasic, respectively.

To extract in an unbiased way the two dominant population dynamics of inhibitory neurons, we applied a two-component non-negative matrix factorisation. One temporal component was positively associated with platform rotation and the other negatively associated with it (Supplementary Fig. 3D-F). Comparing the two components predicted movement state above chance (Supplementary Fig. 3G), indicating that opposing inhibitory population modes carried information about ongoing orienting.

We next classified neurons according to their temporal profile and relative loading onto the two components. Tonic neurons loaded exclusively onto the movement-suppressed component and were designated Tonic^NOGO^. Phasic neurons separated into a movement-coupled group, Phasic^GO^, and a second group with stronger loading on the movement-suppressed component, Phasic^NOGO^. Consistent with this classification, Phasic^GO^ neurons were positively correlated with rotation, whereas Phasic^NOGO^ neurons were only weakly coupled to movement (Supplementary Fig. 3H-J).

A volumetric recording of the SC revealed that Phasic^GO^ and Phasic^NOGO^ neurons occupied partially segregated rostrocaudal bands, while the sparse Tonic^NOGO^ neurons overlapped regions enriched in Phasic^NOGO^ cells (Fig. 1K-M). This observation suggests a spatial organization of movement-related inhibitory dynamics.

Together, these analyses identify three activity profiles within the broader inhibitory population and provide a functional reference for comparing molecularly targeted populations.

### Intersectionally targeted inhibitory populations show distinct biases in movement-related activity

To determine whether molecularly accessible inhibitory populations differed in their movement-related activity, we first examined Sst, Cck and Pvalb expression together with Vgat and Vglut2 by multiplexed FISH. Each marker was present in both inhibitory and excitatory cells, and the fraction coexpressing Vgat varied markedly with marker and SC layer (Supplementary Fig. 4A-F). Nearly all Sst-positive cells in superficial SC were Vgat-positive, whereas the inhibitory fraction was lower in intermediate and deep layers. Cck showed substantially lower Vgat overlap across all layers, and approximately half of Pvalb-positive cells were Vgat-positive. Sst showed little overlap with Cck or Pvalb within the Vgat population, whereas Cck and Pvalb overlapped partially (Fig. 2A-C and Supplementary Fig. 4G-J). Analysis of the Allen Brain Cell Atlas similarly indicated that Sst was enriched in one SC GABA subclass, whereas Cck expression spanned two Pvalb-enriched subclasses (Supplementary Fig. 4K-M)^46^.

**Fig. 2:**
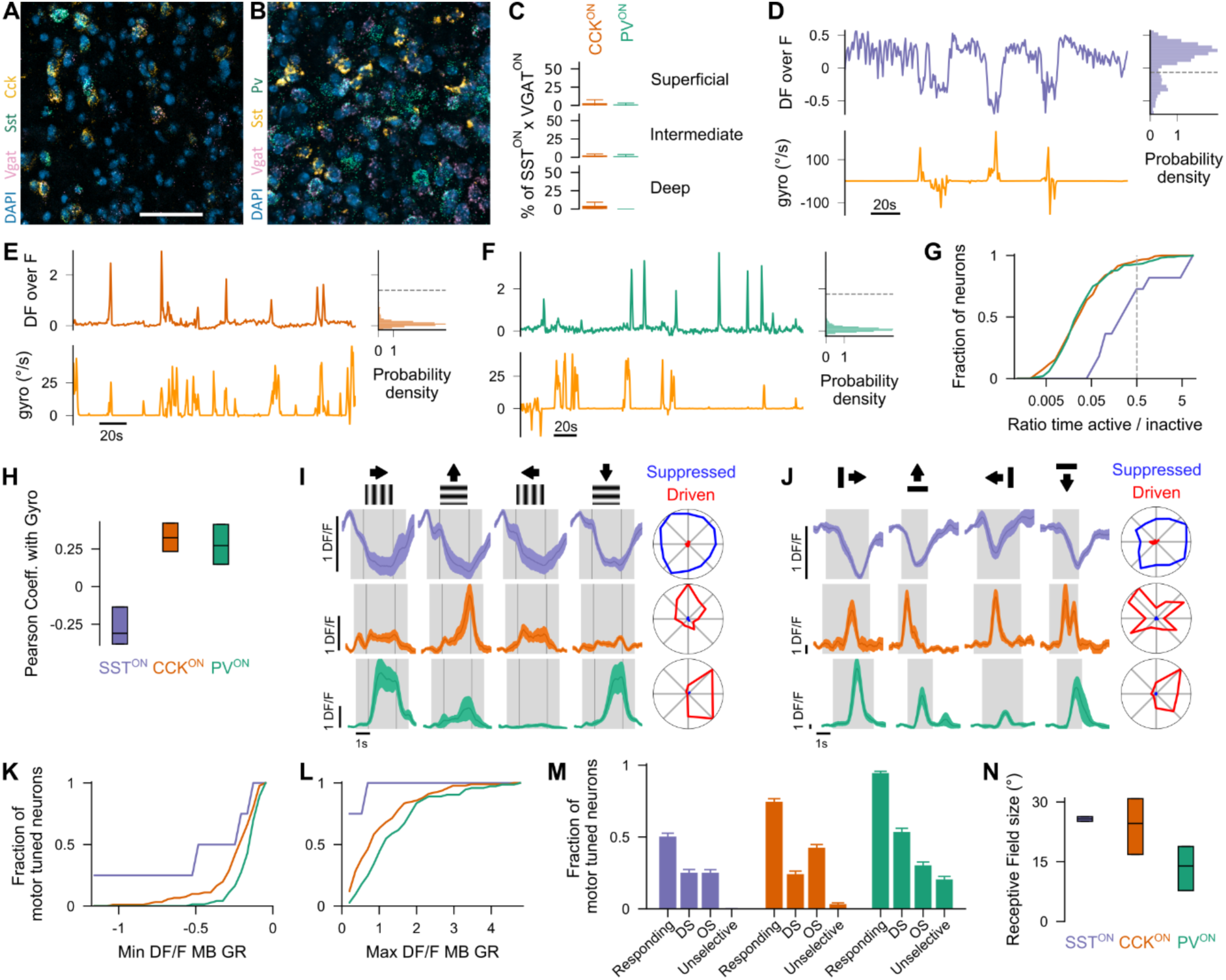
Intersectionally targeted inhibitory populations show distinct movement and visual response biases. **(A)** Example multiplexed FISH image of the SC when probing for Vgat, Sst and Cck. Scale bar represents 50 µm. **(B)** Same as A, but probing for Pv and Sst. **(C)** Mean percentage of neurons coexpressing Cck or Pv together with Sst within the VGAT population for each SC layer. Mean ± standard deviation across three SC coronal sections for the superficial, intermediate and deep layers, respectively. Sst and Cck positive neurons: 2.9 ± 5.0; 2.4 ± 2.1; 4.6 ± 4.8. Sst and Pv positive neurons: 1.4 ± 1.9; 1.5 ± 2.1; 0 ± 0. **(D)** Top: example calcium trace of a SST^ON^ neuron. Bottom: synchronized platform rotation speed. Right: distribution of DF over F timepoints for the recording duration. Y axis is the same as on the left trace. **(E-F)** Same as D for CCK^ON^ and PV^ON^ neurons in e and f, respectively. **(G)** Cumulative distribution of the ratios between the number of DF over F timepoints above and below the midrange value for SST^ON^ (blue), CCK^ON^ (orange) and PV^ON^ (green) neurons. Dashed line represents the threshold used to classify neurons as tonic or phasic. Pvalue for two-tailed two-sample Kolmogorov-Smirnov test; SST-CCK and SST-PV: < 0.0001, PV-CCK: 0.63. **(H)** Pearson correlation coefficients between the platform rotation speed and the calcium traces of neurons. Median [interquartile range]. SST^ON^: −0.31 [−0.38 −0.14]; CCK^ON^: 0.32 [0.23 0.42]; PV^ON^: 0.27 [0.15 0.41]. **(I)** Example calcium responses to drifting gratings for each inhibitory subpopulation. Arrows above indicate the moving direction of gratings. Grey shade indicates the time the grating was on the screen. The left and right thin lines within each grey patch indicate the time when the gratings started and stopped drifting, respectively. Response curves correspond to the mean responses across trials (dark line) surrounded by the standard error of the mean. Right polar histograms indicate the mean suppressed and driven responses for all moving directions. **(J)** Same as I for moving bars. Shaded grey patches indicate when the bars appeared on the screen. **(K)** Cumulative distributions of the minimum DF over F values in response to moving gratings and bars for each neuron within each inhibitory subpopulation. **(L)** Same as K for the maximum DF over F values. **(M)** Fraction of motor tuned neurons responding, DS and OS for either moving bars or gratings. Unselective were the motor tuned neurons which responded to moving bars or gratings and were not DS or OS. Error bars represented the mean ± standard deviation values following bootstrapping. Responding: SST^ON^ 0.50 ± 0.05; CCK^ON^: 0.75 ± 0.04, PV^ON^: 0.94 ± 0.02. DS: SST^ON^ 0.25 ± 0.04; CCK^ON^: 0.24 ± 0.04; PV^ON^ 0.53 ± 0.05. OS: SST^ON^ 0.25 ± 0.04; CCK^ON^ 0.43 ± 0.05; PV^ON^ 0.30 ± 0.05. Unselective: SST^ON^: 0.00 ± 0, CCK^ON^: 0.03 ± 0.02, PV^ON^: 0.21 ± 0.04. **(N)** Receptive field sizes in degree inferred from the delay in response to the moving bars for each subpopulation. Median [interquartile range]: SST^ON^ 25.73 [25.13, 26.33]; CCK^ON^ 24.60 [16.82, 30.85]; PV^ON^ 13.92 [7.76, 18.82]. N_neurons_ = 120.

Because marker expression alone did not isolate inhibitory cells, we used intersectional genetics to target Vgat-expressing cells that also expressed Sst, Cck or Pvalb. We refer to these populations as SST^ON^, CCK^ON^ and PV^ON^, respectively. We recorded their activity during spontaneous orienting in darkness and, in a subset, with the eyes covered (Supplementary Fig. 5A-E). The populations showed clear distinct biases with SST^ON^ were enriched for tonic dynamics and negatively coupled to movement at the population level; CCK^ON^ and PV^ON^ neurons were enriched in phasic dynamics, with CCK^ON^ showing the strongest positive movement coupling (Fig. 2D-H and Supplementary Fig. 5F-I). Only a subset of cells in each population met the motor-tuning criterion (Supplementary Fig. 5J).

A linear model trained on population activity predicted platform rotation above shuffled controls for each targeted population. CCK^ON^ recordings provided the strongest predictions, followed by SST^ON^ and PV^ON^ recordings (Supplementary Fig. 5K-O). This analysis quantifies population-level information about movement and is independent of the NMF-based classification.

Thus, the molecularly targeted populations are not homogeneous functional classes. Rather, SST^ON^ neurons are enriched for tonic, movement-suppressed dynamics relative to CCK^ON^ and PV^ON^ neurons, while CCK^ON^ neurons show the strongest movement-coupled population activity.

### SST and CCK inhibitory populations show opposing stimulus-aligned activity

We next characterized activity during presentation of moving gratings, bars and spots (Supplementary Fig. 6A). Among motor-tuned neurons, SST^ON^ cells tended to decrease activity during visual stimulation, whereas CCK^ON^ and PV^ON^ cells tended to increase activity (Fig. 2I-L). More than half of the motor-tuned cells in each population met the visual-response criterion, and many were direction- or orientation-selective (Fig. 2M)^47–51^. PV^ON^ neurons were enriched for direction-selective responses, whereas CCK^ON^ neurons were enriched for orientation-selective responses; nearby cells tended to have more similar preferences (Supplementary Fig. 6D-G). SST^ON^ and CCK^ON^ neurons also had broader estimated receptive fields than PV^ON^ neurons (Fig. 2H). Together, these results reveal opposing population biases during visual stimulation, with SST^ON^ activity generally suppressed and CCK^ON^ activity generally enhanced.

### SST inhibitory neurons constrain visual recruitment of Pitx2 spatial-motor modules

Because SST^ON^ activity was negatively coupled to orienting and visual stimulation, we asked whether these neurons provide local input to Pitx2^ON^ spatial-motor neurons (Fig. 3A). In a monosynaptic rabies tracing experiment from Pitx2^ON^ neurons, Sst- and Vgat-expressing cells were identified among local presynaptic inputs (Fig. 3B-G). This anatomical observation motivated a direct functional test of whether SST^ON^ population activity constrains Pitx2^ON^ recruitment.

**Fig. 3:**
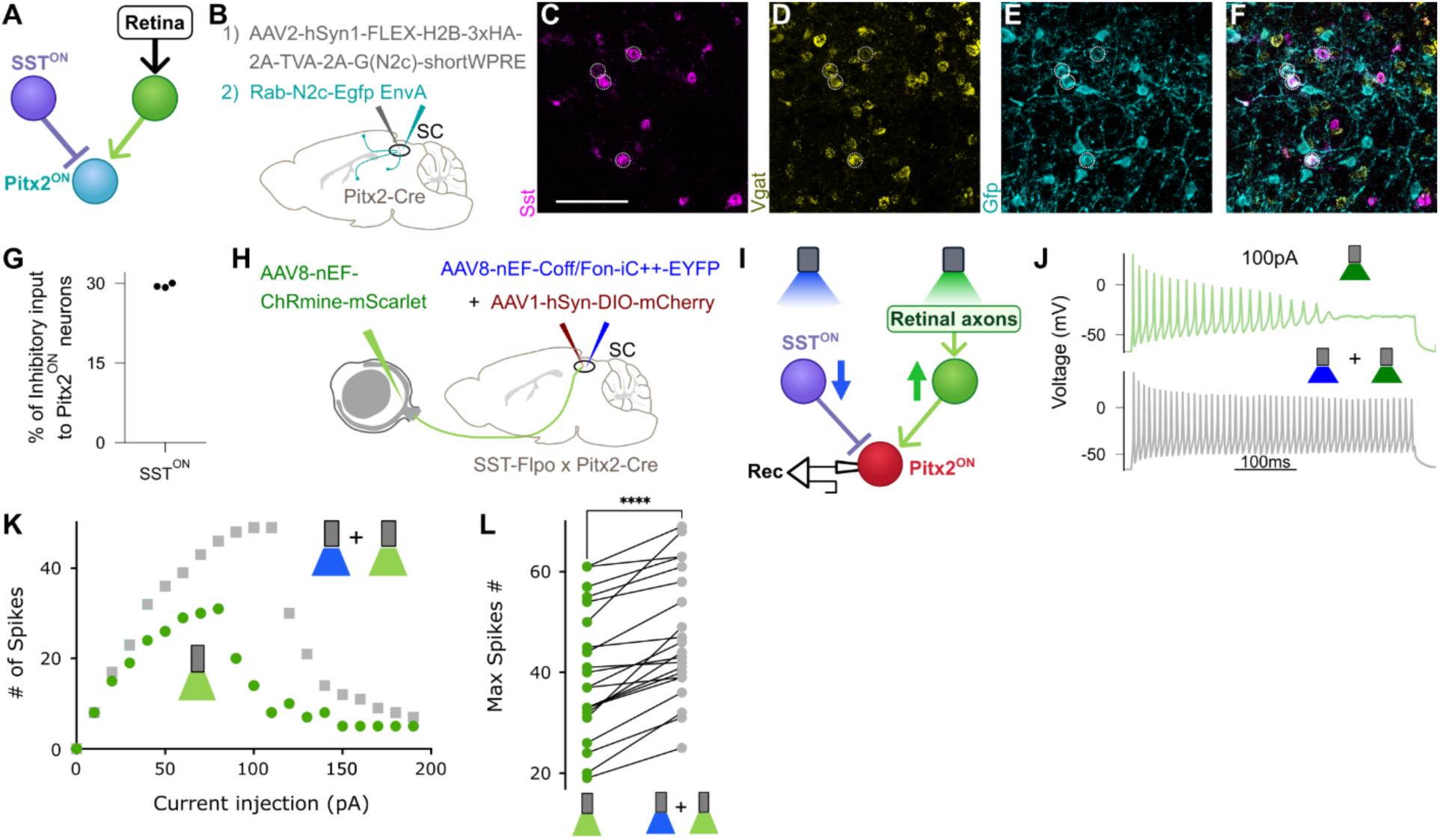
SST neurons modulate visually driven recruitment of collicular spatial-motor modules. **(A)** Model of SST^ON^ control of Pitx2^ON^ visual responses. Activity in the SST^ON^ population is predicted to inhibit Pitx2^ON^ neurons and thereby reduce the likelihood that retinal excitation recruits their firing. The blunt line represents inhibition and the arrow excitation. **(B)** Retrograde monosynaptic rabies strategy to label inputs to Pitx2^ON^ neurons with EGFP. **(C-F)**, Example SC field showing Sst RNA, Vgat RNA, Gfp RNA and the overlay. Dotted circles indicate Sst- and Vgat-expressing cells among EGFP-labelled local inputs. Scale bar, 100 µm. **(G)** Percentage of Sst- and Vgat-expressing cells among local Vgat-expressing inputs to Pitx2^ON^ neurons. Each dot represents a coronal section; mean ± s.d. across three sections spanning the anteroposterior axis, 29.53 ± 0.34%; one mouse. **(H)** Experimental strategy to test whether SST^ON^ population activity constrains Pitx2^ON^ responses. ChRmine was expressed in retinal ganglion cell axons, iC++ in SST^ON^ neurons and mCherry in Pitx2^ON^ neurons. **(I)** Retinal axons were stimulated with green light either alone or together with blue-light silencing of SST^ON^ neurons. **(J)** Example current-clamp responses from a Pitx2^ON^ neuron. **(K)** Number of spikes evoked under the two light conditions across current-injection steps for the same example neuron in J. **(L)** Paired maximum spike counts across recorded neurons. Paired t-test, P < 0.0001; 11 mice, 22 neurons.

We expressed ChRmine in retinal ganglion cell axons, iC++ in SST^ON^ neurons and mCherry in Pitx2^ON^ neurons, then recorded Pitx2^ON^ cells in acute slices (Fig. 3H-I). Retinal axon stimulation evoked Pitx2^ON^ firing, and simultaneous silencing of SST^ON^ neurons increased the number of evoked action potentials across a broad range of current-injection levels (Fig. 3J-L). Thus, activity in the SST population limits the efficacy with which retinal input recruits Pitx2 spatial-motor neurons.

### Increasing SST inhibitory tone reduces responses to lower-contrast visual targets

To test whether changing SST^ON^ activity alters visually guided behaviour, we expressed hM3Dq bilaterally in SST^ON^ neurons and administered either DCZ or PBS. c-Fos labelling was higher after DCZ than after PBS, confirming that the manipulation increased activity in the targeted population (Fig. 4A-F).

**Fig. 4.**
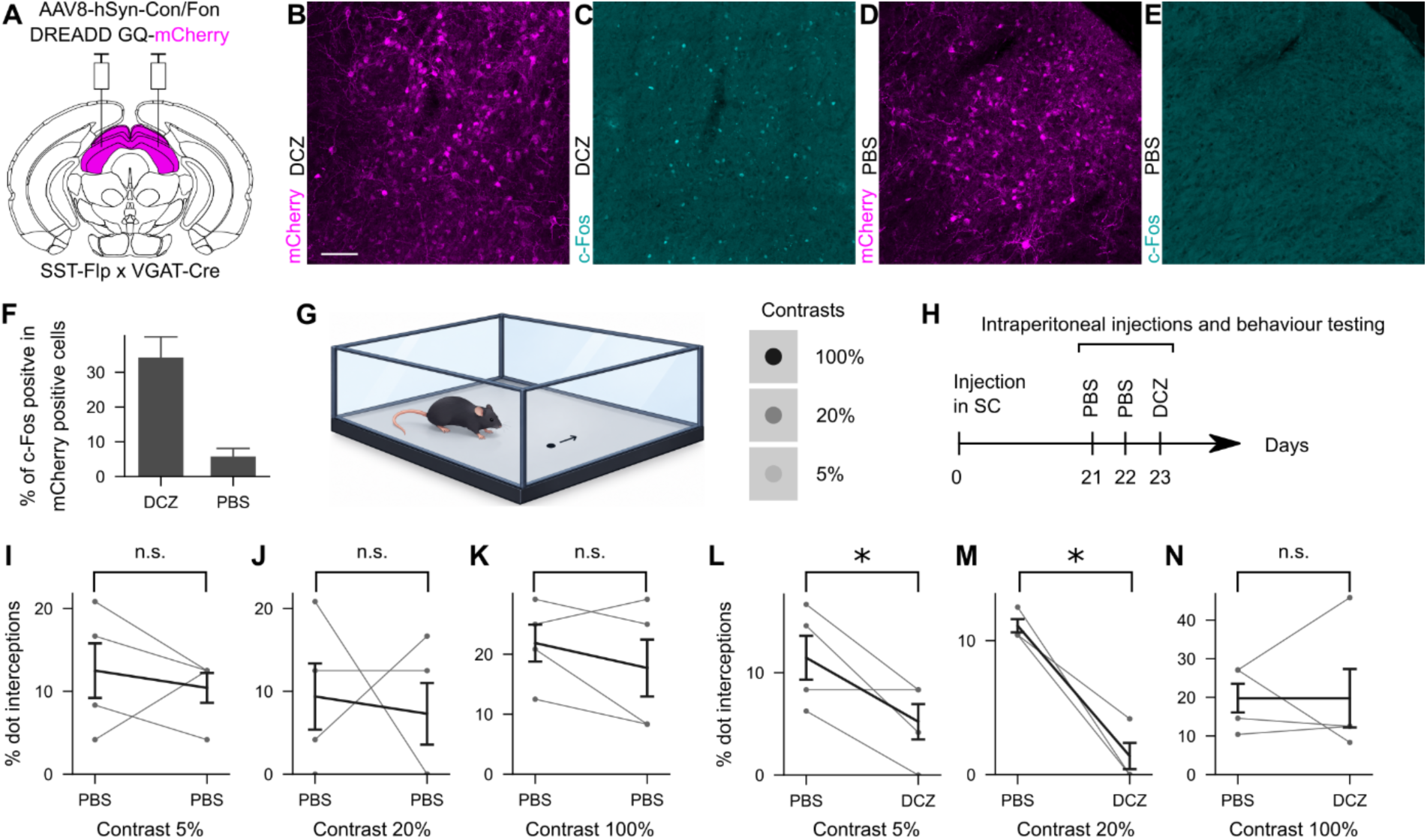
Enhancement of SST activity reduces interception of lower-contrast visual targets. **(A)** AAV injections to label SST^ON^ neurons across both superior colliculi (in magenta) of SST-Flp x VGAT-Cre mice. **(B-C)** Example fields of view of the SC showing the expression of mCherry in B and c-Fos in C following intra-peritoneal injection of DCZ 1h before the mouse was perfused. Scale bar represents 100 µm. **(D-E)** Same as in B-C but administering PBS instead. **(F)** Percentage of c-Fos positive neurons among mCherry positive neurons. Error bars indicate mean ± standard deviation across three field of view spanning the rostro-caudal axis of the SC. DCZ: 34.29 ± 11.86, N_mice_ = 2; PBS: 5.78 ± 4.58, N_mice_ = 2. **(G)** Schematic of the behavioural set up used to test innate orienting of mice towards visual stimuli of different contrast levels. Mice were free to move on an arena with a transparent bottom that stood over a monitor. A single dark dot with three possible contrasts, 5% (low), 20% (middle) and 100% (high), was presented moving in different directions. **(H)** Timeline of the experiment used to test the effect of activating SST^ON^ neurons on the orienting of the mouse. Mice were tested three weeks after the virus was injected in the SC. On the first two days, mice were administered PBS intraperitoneally 20 minutes before entering the orienting arena. The third day, mice were given DCZ instead. **(I-K)** Paired percentages of dot interceptions between the first two days under PBS administration, for each dot contrast (5% in I, 20%, in J and 100% in K). Day 1 and day 2 are on the left and right in each graph, respectively. Thin grey lines indicate individual mice. Black thicker error bars correspond to the average performance across mice. Top indicated the significance of a paired t-test testing whether the performance was greater of day 1 compared to day 2. n.s means non-significant (P value > 0.05) and * means that the P value was below 0.05. Mean ± standard deviation across mice. 5%; Day1: 12.50 ± 6.59, Day2: 10.42 ± 3.61. 20%; Day1: 9.38 ± 8.00, Day2: 7.29 ± 7.44. 100%; Day1: 21.88 ± 6.16, Day2: 17.71 ± 9.49. N_Mice_ = 4. **(L-N)** Same as I-K comparing instead the mean performance of mice for the first two days under PBS with their performance under DCZ on day 3. 5%; PBS: 11.46 ± 4.29; DCZ: 5.21 ± 3.45; 20%; PBS: 11.11 ± 0.98; DCZ: 1.39 ± 1.96; 100%; 19.79 ± 7.44; DCZ: 19.79 ± 15.13; Only mice that intercepted more than 5% of the dots when administered PBS during the first 2 days were kept for the DCZ test on day 3.

We then used a freely moving dot interception assay in which target contrast was varied while position and movement were tracked (Fig. 4G-H and Supplementary Video 2). In four mice tested with PBS during the first two days, DCZ administration on the third day was associated with fewer interceptions of 5% and 20% contrast targets, whereas interception of 100% contrast targets was not detectably changed (Fig. 4I-N). These data provide initial behavioural evidence that increasing SST^ON^ tone preferentially reduces the efficacy of weaker visual stimuli in recruiting target-directed behaviour while not affecting motor execution.

### Local and brain-wide inputs identify candidate regulators of SST inhibitory neurons

We next asked which inputs could regulate the SST^ON^ population. Because CCK^ON^ neurons showed movement- and stimulus-related activity opposite to SST^ON^ neurons, we tested whether they were among local presynaptic inputs. Monosynaptic rabies tracing from SST^ON^ neurons identified Cck- and Vgat-expressing cells within the SC (Fig. 5A-F and Supplementary Fig. 7A-C). This result provides an anatomical substrate for a candidate local inhibitory control of SST^ON^ neurons via CCK^ON^ neurons.

**Fig. 5.**
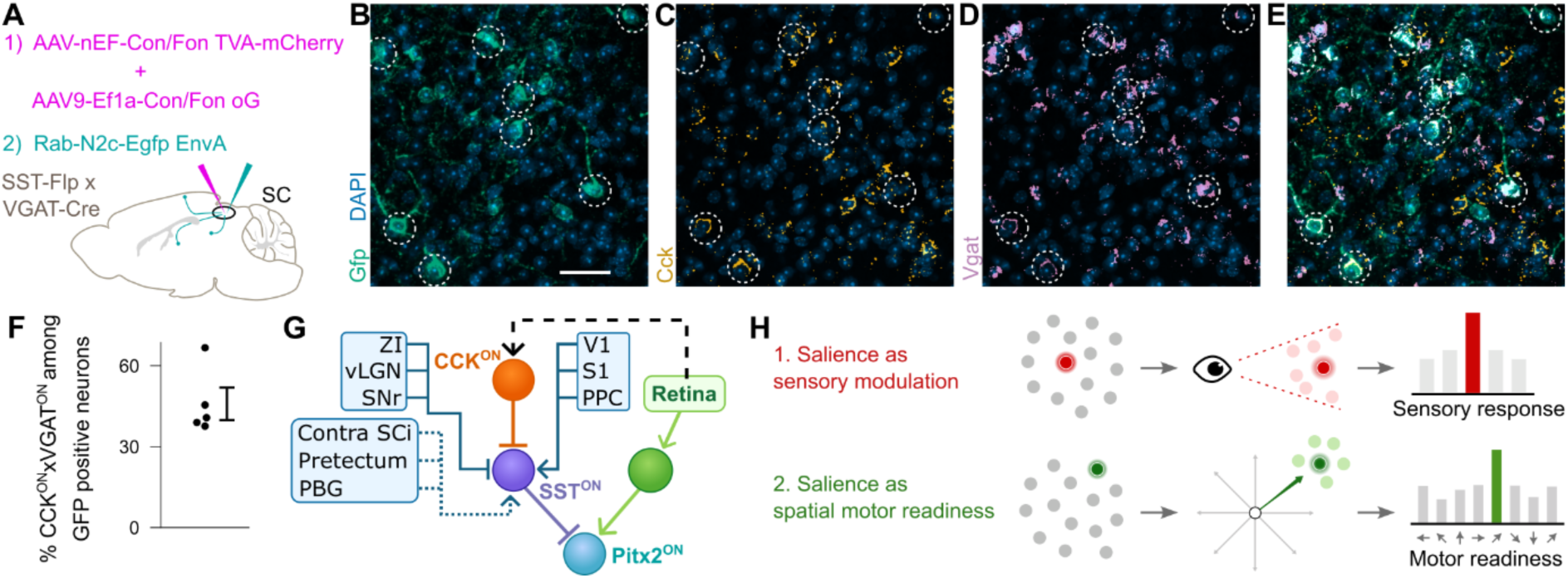
Local and extra-collicular inputs identify candidate routes for regulation of the SST gate. **(A)** Monosynaptic retrograde rabies strategy to label inputs to SST^ON^ neurons. Rabies virus was injected three weeks after the helper AAVs. **(B-E)** Example SC field showing Gfp RNA, Cck RNA, Vgat RNA and the overlay. Dotted circles indicate Cck- and Vgat-expressing cells among labelled local cells. Scale bar, 50 µm. **(F)** Percentage of Cck- and Vgat-expressing cells among GFP-positive cells in five coronal sections; mean ± s.d., 45.89 ± 11.99%; one mouse. Because GFP-positive cells included both starter cells and presynaptic inputs, this value should be interpreted as a descriptive estimate rather than an input fraction. **(G)** Proposed circuit model. Local CCK^ON^ neurons and extra-collicular afferents are anatomically positioned to regulate SST^ON^ activity and thereby influence SST-dependent control of Pitx2^ON^ visual recruitment. The signs and behavioural functions of the extra-collicular pathways, and the functional CCK^ON^-to-SST^ON^ interaction, remain to be tested. Abbreviations are listed in Supplementary Fig. 7. **(H)** Two conceptual mechanisms by which salience can influence behaviour. Top, in a sensory-modulation framework, salient stimuli are preferentially processed through enhancement of their sensory representation. Bottom, in a spatial motor-readiness framework, salient stimuli preferentially increase the readiness of the corresponding motor module to be recruited for orienting.

Brain-wide tracing also identified inputs from the zona incerta, ventral lateral geniculate nucleus, substantia nigra pars reticulata, visual cortex, somatosensory cortex, posterior parietal cortex, parabigeminal nucleus, pretectum and contralateral SC (Fig. 5G and Supplementary Fig. 7D-L). Based on the known transmitter phenotypes of these regions, some pathways are positioned to inhibit SST^ON^ neurons whereas cortical pathways are positioned to excite them. These projections therefore identify candidate routes through which sensory, motor and contextual signals could regulate the SST gate.

## Discussion

Here we identify a somatostatin-expressing inhibitory population as a control point between retinal input and collicular motor output. SST^ON^ neurons showed a population bias toward tonic, movement-suppressed activity and tended to reduce their activity during visual stimulation. Silencing this population increased retinally evoked firing in Pitx2^ON^ spatial-motor neurons, establishing that somatostatinergic inhibition constrains the recruitment of collicular output by visual input. The central advance is therefore not simply that inhibition influences SC function, but that a molecularly accessible inhibitory population controls the input-output relationship of a genetically defined spatial-motor pathway.

The behavioural manipulation is consistent with this interpretation. Increasing SST^ON^ tone reduced interception of lower-contrast targets while sparing responses to the highest contrast tested. Together with the slice physiology, this pattern supports a graded gate in which somatostatinergic inhibition changes the probability that a visual signal recruits orienting-related output. Here, “gate” refers to regulation of sensory-to-motor transfer, not an absolute block on movement or a uniquely localized action threshold.

These findings suggest a complementary view of salience. Rather than requiring salient stimuli to be represented more strongly only within sensory pathways, their behavioural impact can also depend on the state of the premotor circuits receiving those signals (Fig. 5H). In this framework, SST-dependent inhibition regulates the readiness of spatial-motor modules for recruitment, such that the same sensory input can have different behavioural efficacy depending on premotor state. We therefore do not propose that SST neurons encode salience as a sensory variable; rather, they provide a mechanism through which salience can acquire differential access to action.

The movement-related suppression of SST^ON^ activity indicates that the gate is not exclusively visual. Signals related to locomotor, postural or internal state could lower somatostatinergic tone during ongoing behaviour and thereby alter the gain of retinal input^52^. Such state dependence offers a parsimonious explanation for why the same visual event can have different behavioural efficacy across behavioural contexts. It also distinguishes SST-mediated control from a final motor brake: increasing SST^ON^ activity did not abolish responses to high-contrast targets, suggesting modulation of recruitment rather than an obligatory permissive step for movement.

The input mapping identifies possible routes by which this control could be adjusted. Local CCK^ON^ neurons are anatomically positioned to inhibit SST^ON^ neurons and show an opposing population response during visual stimulation, together supporting a candidate disinhibitory motif. Likewise, cortical and subcortical inputs provide an anatomical substrate for contextual regulation, but the current tracing data do not assign their sign, behavioural role or effect on SST^ON^ activity.

Because Pitx2^ON^ neurons are organized into spatial-motor modules^9,15^, local regulation of their SST-mediated inhibition could in principle make sensory access spatially selective. A focal decrease in SST^ON^ activity could increase the responsiveness of a corresponding motor module, whereas focal increases could reduce recruitment of that region of the collicular map.

Together, our results establish that premotor inhibition can regulate the efficacy with which visual input recruits spatially organized motor output. They identify the SST^ON^ population as a causal control point and reveal a broader anatomical architecture through which local and extra-collicular signals could regulate that control. These findings extend sensory accounts of salience by identifying the motor readiness of spatial-motor circuits as an additional determinant of whether sensory information gains behavioural access to action.

## METHODS

### Mice and surgical procedures

#### Mice

All animal procedures were conducted in accordance with the UK Animals (Scientific procedures) Act 1986 under project license PPL PP0188140 approved by The Animal Welfare and Ethical Review Body (AWERB) committee of the MRC Laboratory of Molecular Biology.

Adult mice of around 2 to 4 months old were used of the following lines: C57BL/6 wild-type (WT), VGAT-Cre (MGI:5141270)^53^, VGAT-Flp (MGI:5806634)^32^, VGlut2-Cre (MGI:5141269)^53^,

VGAT-Cre x GCaMP6f (VGAT-Cre crossed with Rosa-LSL-GCaMP6f, MGI:104735^54^), Pitx2-CRE^55^, VGAT-Cre x SST-Flp (VGAT-Cre crossed with SST-Flp, MGI:5700394^56^, VGAT-Flp x CCK-Cre (VGAT-Flp crossed with CCK-Cre, MGI:88297^32^, VGAT-Flp x PV-Cre (VGAT-Flp crossed with PV-Cre, MGI:3590684^57^, Pitx2-CRE x SST-Cre. Pitx2-Cre mice were screened for the absence of eye and tooth defects before use, consistent with previous studies^9,15,58^.

Animals were housed in 12 hours light/dark cycle with food and water ad libitum.

#### Viruses

AAVs were ordered from Addgene or made in house. Rabies viruses were made in house. AAVs were injected with a final titre of around 1-6 x 10^12^. Rabies viruses were injected with a titre of 2×10^8^. Around 200-300nl of virus was delivered at each injection site except for eye injections. Viruses were the following: AAV2/9::CAG-GCaMP6s-WPRE (#100844, Addgene), AAV2/9-flex-GCaMP6s (#100842, Addgene), AAV5-pAAV-EF1a-Con/Fon-GCaMP6f (#137122, Addgene)^59^, AAV8-pAAV-nEF-ChRmine-mScarlet (#137158, Addgene), AAV8-pAAV-nEF-Coff/Fon-iC++-EYFP (#137157, Addgene), AAV1-pAAV-hSyn-DIO-mCherry (#50459, Addgene), AAV2-hSyn1-FLEX-H2B-3xHA-2A-TVA-2A-G(N2c)-shortWPRE^9^, Rab-N2c-Egfp-EnvA^9^, AAV8-hSyn-con/fon-DREADD-Gq-mCherry (#200661, Addgene), AAV9-nEF-Con/Fon-TVA-mCherry (#131779, Addgene), AAV9-Ef1a-Con/Fon-oG (#131778, Addgene)

#### Surgeries

Mice were anaesthetized with isoflurane delivered at a flow of 5% in 2 L/min of O_2_ for the initial induction and then maintained at 1%–2% in 2 L/min of O_2_ for the duration of the surgery. Mice were administered subcutaneously Rimadyl as a pain relief (2 mg/kg body weight) shortly after the induction and before any procedure was carried out. For the cranial window surgery, mice were administered Dexafort at 2 μg/g on the day prior to the surgery to avoid brain swelling when opening the skull, and as result received Vetegersic at 0.1mg/kg instead of Rimadyl as a pain relief. Mepivicaine diluted 1 in 50 was splashed over the skull as a local anaesthetic before any drilling took place. Mice usually recovered within 15 minutes after the surgery and were placed overnight on a heated cabinet supplied with food mash and hydrogel. Their weight was measured for 7 days post-surgery.

#### Brain Injections

Mice were placed on a stereotaxic frame. Their skin was opened, and a small hole was drilled onto the skull. A glass pipette filled with virus pierced into the brain and a Nanoject (Scientific Laboratory Supplies) delivered the virus at a rate of 5nL every 5s. 10 minutes after, the glass pipette was retracted and the skin sutured back together.

#### Miniendoscope implantation

A 1 mm diameter craniotomy and durotomy was performed before the virus was injected into the motor domain of the SC (AP: −3.80 mm, ML: −0.80 mm, DV: −1.70 mm). A GRIN lens (0.5 mm diameter, 3.4 mm length; DORIC Lenses Inc.) was lowered at the same coordinates at a rate of 0.1 mm/min. The exposed brain surface was covered with sterile Vaseline, and the implant was secured with two skull anchor screws and RelyX™ Unicem 2 Self-Adhesive Resin Cement (3M, Bracknell, UK).

#### Cranial window for two-photon imaging

A headpost and a cannula coverslip on top of the rostral left SC were attached to the mice as _in47_

#### Eye injections

Proxymetacaine 0.5% was applied onto the eye for 5 minutes prior to the injection. A hole was pierced behind the lens with a needle while the eye was held with forceps. A glass pipette was inserted into the eye until it touched the retina and then slightly retracted. 1µl of virus was delivered at a rate of 5nl every 2s. Chloramphenicol drops were then applied during the recovery of the mouse.

#### Freely moving calcium imaging recordings

We recorded the head over body positions as in^9^, while neural activity was recorded using a head-mounted miniature fluorescence microscope (DORIC Lenses Inc., Québec, Canada; 2.2 g, 700 × 700 μm field of view, 630 × 630 pixel resolution, 458/35 nm excitation filter, 0.2-2 mW LED intensity) connected to the Doric Neuroscience Studio acquisition system.

Behavioural experiments were conducted in a square Perspex arena (50 × 50 cm) equipped with a lateral-mounted camera. Mice were first handled for one week before undergoing three days of habituation to the arena, the miniscope and the dual body sensors. All behavioural sessions were conducted during the afternoon at a consistent time. Recording consisted of four 10-min trials, two performed under illuminated conditions and two in complete darkness.

#### Head-fixed behaviour

A circular platform with transparent walls floated on a cushion of air. An inertial measurement unit (Arduino Nano 33 BLE) was attached onto the bottom of the platform and communicated its rotation speed via Bluetooth signal.

Mice were comfortable to move about on the platform while being head-fixed after 2 weeks of training. First, mice were handled for a few days before being introduced and head-fixed for brief periods of time on the platform. Mice were then head-fixed for increasing durations until they underwent comfortably 45 minutes sessions.

In a subset of experiments, we covered the eyes of the fully trained mice using loose black tape.

For training mice to capture crickets on the floating platform while being head-fixed, we first food deprived them and started introducing crickets in their home cage as part of their diet.

When mice were habituated to the floating platform as described above and drawn to capture crickets in their home cages, we then only offered crickets to the mice when they were head-fixed on the platform.

### Two-photon calcium imaging

#### Microscope

We recorded the calcium activity of neurons in the SC of head-fixed mice as in^9^.

For the volumetric recording, successive planes separated by SC depth of 20 to 50µm where imaged quasi-simultaneously one after the other by rapidly moving the objective up and down using a piezo device (PFM450E, Thorlabs). The resulting volume recording rate ranged from 1 to 4 Hz.

#### Processing of two-photon recordings

Two-photon recordings were processed as in^9^ with Python and CaImAn^60^ (Flatiron Institute).

#### Traces extraction

Raw calcium traces corresponded to the average pixel intensity values comprising the cell’s region of interest for every timepoint. Raw calcium traces were first convolved with a Hanning window of 1.5 seconds wide, before being detrended. The normalised calcium fluorescence variations (DF over F) were computed by dividing and offsetting the traces with their the 20^th^ percentile value.

Gyroscope traces were also similarly convolved and offsetted by the median value of the non-movement period to remove any drift.

### Analysis of neurons’ motor related properties

#### Responding neurons to spontaneous orienting

For each timepoint, we considered a baseline and a response period of 10 seconds prior and 3 seconds post, respectively. The timepoint’s z-score was: 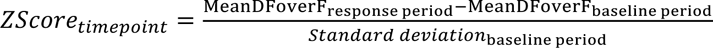. Finally, the neuron’s z-score was the maximum of timepoints’ z-score for the duration of the recording. Neurons were considered to respond if their z-score was above 5.

#### Classification of tonic and phasic neurons

We computed the ratio between the number of timepoints with a DF over F value greater and lower than the midrange. Tonic and phasic neurons had a ratio above and below 0.5, respectively.

#### Non-negative Matrix Factorisation (NMF)

We first rectified the negative DF over F values by changing them to 0. We then used the NMF algorithm from the Python scikit-learn library as in^61^

#### Prediction of mouse movement using NMF temporal components

For each timepoint, we compared W^GO^ with W^NOGO^. If W^GO^ > W^NOGO^, the mouse was inferred to be moving. To test the movement predictions, we defined movement periods when the rotation speed of the platform was above 5 degree per second. We then used a stratified k fold cross validation strategy to divide the data into 3 parts containing equal amounts of moving and no moving periods. The NMF was fitted using 2 parts and the prediction was tested on the remaining third. This process was repeated with three random partitions 10000 times. As a control, we circularly delayed each neuron’s DF over F by a random duration.

#### Motor tuned neurons

Motor tuned neurons were responding neurons with an absolute Pearson correlation coefficient with the platform rotation speed above 0.1 and a statistically significant p value below 0.05, tested by circularly shifting the traces.

We used a bootstrapping analysis to assess the percentages of motor tuned neurons. This consisted in randomly choosing with replacement 20 neurons from each subpopulation and computing the percentages of motor tuned neurons. We repeated this process 10000 times and reported the mean and standard deviation across all repeats.

#### Prediction of movement using a linear model

We predicted the platform rotation speed using a linear model (scikit-learn’s linear regression) fitted on neuronal activity. The data was split in three parts using a stratified K fold strategy and a cross validated mean R^2^ score was computed. We repeated this process 1000 times with random data partitions. For the control, we used circularly delayed DF over F neuronal traces.

#### RNA – In situ hybridization (FISH)

12 μm-thick cryosections were obtained from fixed and frozen brains. Using a cryostat (Leica CM1950) and processed for RNA in situ detection using the RNAscope™ Multiplex Fluorescent V2 Assay according to the manufacturer’s instructions (Advanced Cell Diagnostics, Hayward, California, USA). RNAscope® probes were used to detect mRNA expression of Vgat (gene Slc32a1 #319191), Vglut2 (gene Slc17a6 #319171-C3), Pv (gene Pvalb #421931-C2/C3), Sst (gene Sst #404631-C2/C3) and Cck (gene Cck, #402271-C2/C3). Fluorophores used for detection of mRNA expression were Alexa Fluor 488, Atto 550, Atto 647; TSA Vivid™ 520, 570, and 650 (Tocris 323271, 323272, 323273); 4′,6-diamidino-2-phenylindole (DAPI) was used for counterstaining.

### Visual stimulation

#### Set up

Visual stimuli were presented to the mouse on a grayscale curved LCD monitor (Samsung C27F398FWU, 10 Lux mean luminance, 60 Hz refresh rate), controlled by a custom software written in Octave using Psychtoolbox. The monitor was 40 cm apart and covered a visual field of 73° azimuth and 46° elevation contralateral to the imaged SC. Gray values were gamma corrected to yield linearly perceived changes in brightness.

#### Visual stimuli

A series of full field drifting gratings and moving bars were presented moving into one of 12 angular directions sampling the 360° wheel with 30° increments. Gratings and bars were presented in separated blocks. The sequence of angle presentation was random. Each stimulus was repeated 5 times with an inter-stimuli time gap that varied between 2 and 3 seconds.

Gratings had a spatial frequency of 0.125 cycles per degree and a Michelson contrast of 1. They stayed static for 1 second after appearing on the screen, then drifted with a temporal frequency of 3.2 Hz for 2.5 seconds and finally remained static for another second before the screen returned to a grey background.

Bars were black, 8° wide and swept a grey screen at 40° per second.

In addition, we presented single black spots moving on a grey background in a subset of experiments. Spots were 8° in diameter and moved at a speed of 40° per second alongside the vertical or horizontal axes. For each axis, we moved the spots along three parallel lines, to sample the azimuth and elevation extent of the screen. Therefore, we presented spots moving along a total of 12 trajectories (3 parallel vertical and horizontal lines x 2 opposite directions per line). For each neuron, we considered the trajectories that yielded the highest responses per direction. These were the cardinal trajectories that best overlapped with the receptive field of the neuron.

#### Driven versus suppressed responses

Driven responses consisted in positive variation in calcium activity upon stimulus presentation whereas suppressed responses consisted in negative variations.

#### Quality index

The quality index was computed as in^62^. Here, we set the threshold for responding neurons to 0.3.

#### DS and OS neurons

Neurons responding to visual stimuli were classified as DS or OS as in^47^. Here, DS and OS neurons were not mutually exclusive, so DS neurons could also be OS and vice-versa.

#### Bootstrap analysis to quantify percentage of responding, DS, OS and unselective neurons

We randomly selected with repetition a sample of 100 neurons from each subpopulation’s motor tuned neurons. We then computed the percentage of responding, DS, OS and unselective neurons within this sample. We repeated this 10000 times.

#### Whole-cell patch-clamp recordings and optogenetic manipulation

Acute brain slices were prepared in ice-cold sucrose-based cutting solution containing the following, in mM: 206 sucrose, 2.5 KCl, 1.25 NaH₂PO₄, 26 NaHCO₃, 25 glucose, 5 MgCl₂, and 1 CaCl₂. The cutting solution was continuously bubbled with 95% O₂ / 5% CO₂. After slicing, slices were transferred to ACSF containing the following, in mM: 119 NaCl, 2.5 KCl, 1.25 NaH₂PO₄, 26 NaHCO₃, 20 glucose, 1 MgCl₂, and 2 CaCl₂. ACSF was continuously oxygenated with 95% O₂ / 5% CO₂ during slice recovery and electrophysiological recordings.

Patch pipettes were filled with an intracellular solution containing in mM: 120 K-gluconate, 9 KCl, 3.5 MgCl₂, 4 NaCl, 10 HEPES, 4 Na₂ATP, 0.4 Na₃GTP, 0.5 CaCl₂, and 0.3 EGTA. The pH was 7.2–7.3, and the osmolarity was 270–285 mOsm.

Electrophysiological signals were acquired using the pCLAMP software suite and analyzed offline using Clampfit, Molecular Devices. After establishing the whole-cell configuration, recording quality was first assessed in voltage-clamp mode using test pulses. Cells were included for further analysis only if the input resistance was greater than 300 MΩ, the series resistance was less than 25 MΩ, and these parameters changed by less than 20% during the recording.

After the quality-control assessment, recordings were switched to current-clamp mode. The membrane potential was maintained at −75 mV by injecting bias current when necessary. Neuronal excitability was examined by injecting 500-ms depolarizing current steps starting at 0 pA and increasing in 10-pA increments.

For optogenetic manipulation, green or blue light (pE-300, CoolLED) was shone onto the sample during the 500-ms current-injection periods.

#### Immunohistochemistry and analysis

Histology was performed as in^9^. Primary antibodies were: Goat-anti-RFP (Antibodies.com, A121675, 1:1000), Rabbit-anti-c-fos (Cell Signaling, 2250S, 1:500), Chicken-anti-GFP (Aves, GFP-1020, 1:2000). Secondary antibodies were diluted 1 in 1000: Alexa Fluor™ 488 donkey-anti-rabbit (Molecular Probes, A21206), Alexa Fluor™ 555 Donkey-anti-Goat (Invitrogen, A-21432), Alexa Fluor 488™ donkey-anti-chicken (Invitrogen, A78948).

Images were acquired with a confocal microscope (LSM880, Zeiss). Coronal sections were aligned to the Allen Brain Reference Atlas^63^ using QuickNII^64^, DeepSlice^65^ and VisuAlign^66^. GFP or RFP positive cells were segmented using CellPose^67^. Segmented cells were considered positive with respect to another marker if the maximum fluorescent value inside a cell’s outline on the maker’s channel was greater than the mean plus 3 times the standard deviation of whole marker image.

### Behaviour for testing the probability to orient towards stimuli of different contrast

#### Behaviour set up

The arena was made of 4 red transparent Perspex^TM^ sheets that formed the walls and of a clear cast Perspex^TM^ on the bottom. The base was a rectangle of 600mm x 330mm size and the walls were 300mm high.

The arena stood over a computer monitor (Dell U2715H) where we presented single moving dots of 8mm in diameter cruising at a constant speed of 40mm per second over a grey background (mean illuminance 13 lux). Gray values were gamma corrected to yield linearly perceived changes in brightness. We presented dots moving in 8 possible directions sampling the 360° wheel with 45° intervals, and 4 of these angles corresponded to the symmetrical axis of the arena. Dots had either 5, 20 or 100% contrast defined by the inverted Webber score: 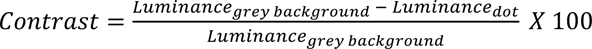. Each combination of dot contrast and moving direction was presented 3 times. Therefore, each contrast dot was presented 24 times to the mouse during an experimental session. A random inter-trial duration of between 6 and 10 seconds separated each dot presentation.

A camera (FLIR, BFS-U3-13Y3M-C) placed above the arena and equipped with a varifocal imaging lens (Edmund Optics, #58-365) captured images at 70Hz via SpinView (FLIR, Spinnaker SDK). The arena was located in a closed enclosure with no lights, except for the brightness of the display monitor. Dot presentations and camera images were synchronised with TTL pulses

#### Analysis

We used DeepLabCut^68,69^ to extract the positions of the nose and head of the mouse in camera coordinates and dot’s instantaneous positions on the screen were also converted into camera coordinates. For each timepoint the distance between the nose of the mouse and the dot was computed and the mouse was considered to have intercepted a dot when the distance between its nose and the dot was less than 12mm.

#### Experimental paradigm

Mice received intra-peritoneal injections of either PBS or DCZ (0.5mg/kg, Hello Bio, HB9126) 20 minutes prior to entering the arena. We considered only mice that intercepted at least 5 % of the dots during the first 2 days with PBS. We then compared the mean percentage of interceptions obtained during the first two days with the percentage of interception achieved under DCZ administration on the third day.

#### Allen Brain Cell atlas analysis of SC GABAergic neurons

We queried the Allen Brain Cell atlas and its Whole-Mouse-Brain (WMB) taxonomy^46^ through the abc_atlas_access Python API. From its spatial (MERFISH) datasets, we selected GABAergic neurons (neurotransmitter “GABA”, excluding non-neuronal supertypes) and localized them to the superficial and motor SC by mapping each cell’s Common Coordinate Framework (CCFv3) coordinates. Retaining the WMB subclasses each representing ≥1% of this SC GABAergic pool yielded 1 class, 9 subclasses and 33 supertypes (Supplementary Fig. 4k), whose spatial distributions were plotted on the section containing the most SC GABA cells (Supplementary Fig. 4l). To examine marker expression at the single-cell level, the same supertypes were retrieved from the WMB 10x single-nucleus RNA-sequencing dataset and analysed for Sst, Cck and Pv mRNA as scanpy dot plots (per-gene-scaled mean expression and fraction of expressing cells) grouped by subclass (Supplementary Fig. 4m).

## Acknowledgements

We thank the Laboratory of Molecular Biology (LMB) electronics and mechanical workshops for the help with hardware development and members of the Biological Service Group for their support with animal husbandry. We thank Ernesto Ciabatti for providing the viruses used in the rabies tracing experiments and Nicolas Alexandre for supporting the analysis of the 1P miniscope data.

## Author contributions

MT and DdM conceptualised the project, designed the experiments and wrote the manuscript. DdM performed all 2P imaging, tracing and behavioural experiments. LM performed the 1P imaging experiments. YZ performed the whole cell recordings in slices. AF performed the *in situ* hybridization experiments. DW and LG provided support with tracing, immunohistochemistry and *in situ* hybridization experiments. FM made all the viral preparations.

## Additional information

### Supplementary figure/table legends

**Supplementary Fig. 1:**
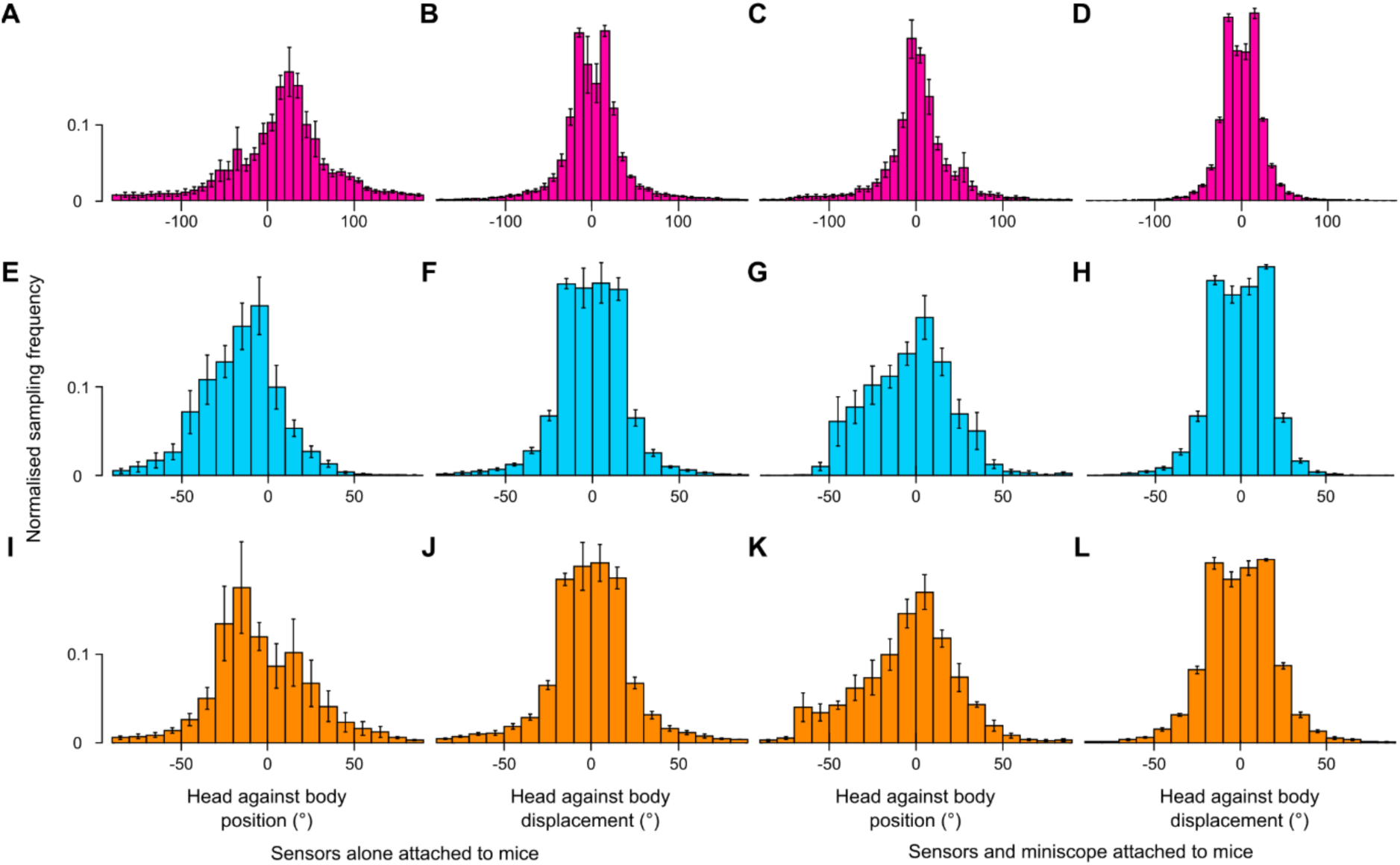
Head mounted miniendoscope did not affect the nature of head rotations. **(A-B)** Distribution of head yaw angle positions in A and displacements in B when mice had attached only the head and body sensors. **(C-D)** Same as A-B when mice had attached the miniendoscope in addition to the body and head sensors. **(E-H)** Same as A-D for the pitch component of the head rotations. **(I-L)** Same as A-B for the roll component of the head rotations. Mice implanted with a miniendoscope exhibited normal head movements, with movement ranges comparable to those observed in mice carrying only the sensors. The distributions of conjunctive movements were not significantly different between the two groups, indicating that miniendoscope implantation did not measurably alter natural movement dynamics. Data are presented as mean ± standard error of the mean across mice.

**Supplementary Fig. 2:**
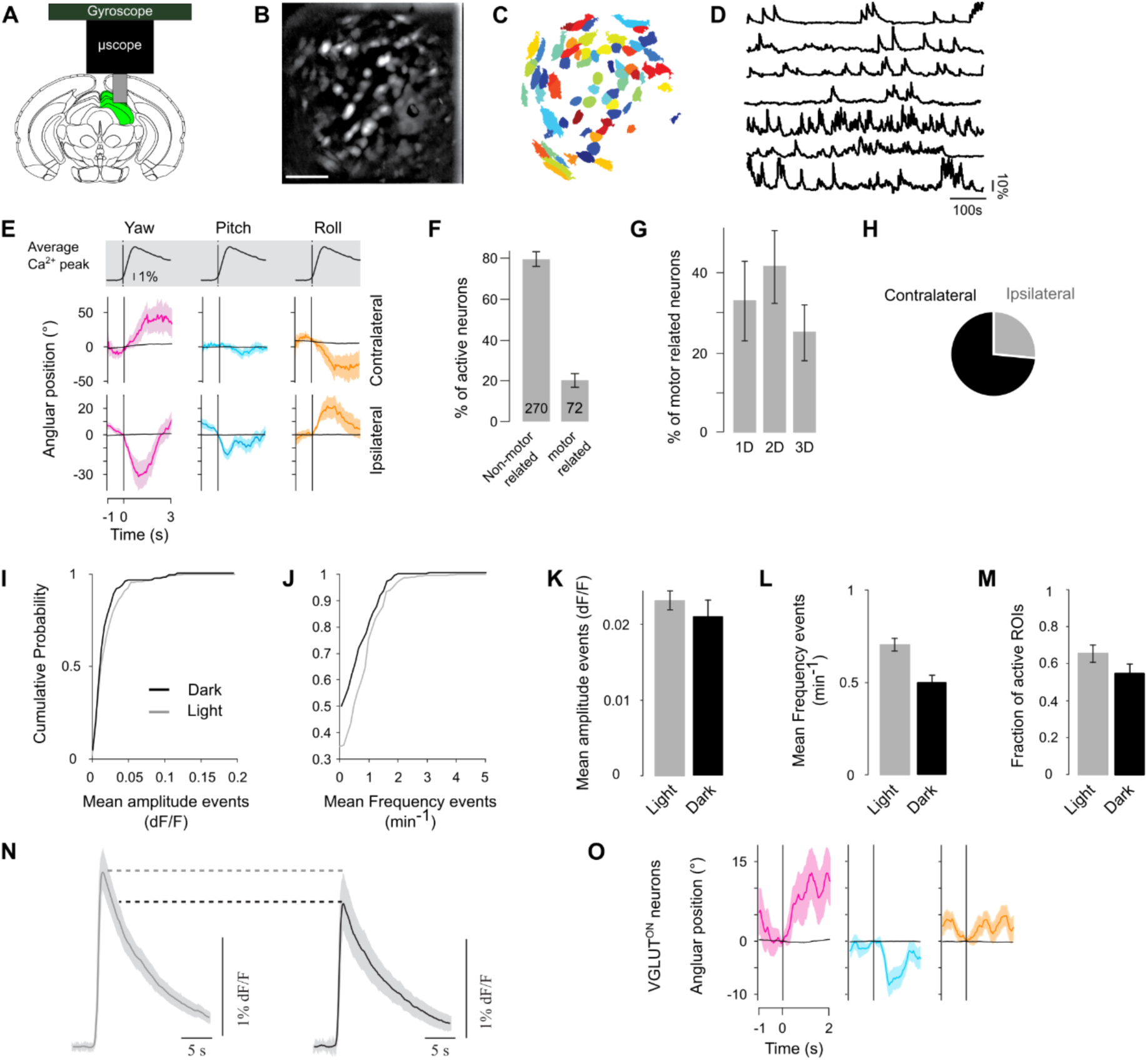
Validation of endoscopic imaging to characterise head orienting movements. **(A)** Cells in the SC (in green) expressed a calcium indicator and their fluorescent activity was recorded through a GRIN lens (grey cylinder) coupled with a microendoscope connected to a gyroscope to simultaneously record head displacements. **(B)** Maximal projection of a time series imaged in the deep layers of the SC. The field of view is 320 µm. Neurons expressing GCaMP6s are visible as white rounded blobs. Neurons here were labelled using pan neuronal GCaMP6s. Scale bar represents 100µm. **(C)** Segmentation of the active region of interests from the same field of view showed in B. **(D)** Representative calcium traces from SC neurons shown in D. **(E)** Example of two neurons’ head tuning curve. Top panel: Average calcium event calculated across all neurons’ calcium events. Shaded area represents the standard error of the mean. Bottom panels: spike triggered average of head displacement along yaw (magenta), pitch (cyan), roll (orange). Spikes were inferred from the deconvolved calcium traces. Shaded areas represent the standard error of the mean. Black curves correspond to the spike triggered average obtained after shuffling the spike times. The vertical lines at time 0 represent the onset of the neuron’s firing events. The top cell was tuned to contralateral yaw rotations, while the bottom cell was tuned to ipsilateral yaw rotations. **(F)** Percentage of non-motor and motor related neurons recorded. Error bars represent mean ± standard error of the mean values. Motor related: 20.24 ± 3.32. **(G)** Percentage of motor related neurons tuned to one, two or three dimensions of head rotations. Error bars represent mean ± standard error of the mean. 1d: 32.99 ± 10.02; 2d: 41.49 ± 9.07; 3d: 25.00 ± 7.04. **(H)** Pie chart showing the fraction of collicular motor related cells tuned to ipsilateral versus contralateral yaw head rotations with respect to the collicular hemisphere of the recording. Percentages are 26.36 ± 4.96 for ipsilateral and 73.12 ± 5.26 for contralateral, Mean ± standard error of the mean. N_mice_=7; N_neurons_=344; N_non motor-tuned_=270; N_motor-tuned_=90. **i**, Cumulative distribution of mean DF over F amplitude of calcium events between light (grey) and dark (black) conditions across neurons. **(J)** Cumulative distribution of mean frequency of calcium events between light (grey) and dark (black) conditions across neurons. **(K)** Mean ± standard deviation of amplitude events across all neurons between light (grey) and dark (black) conditions. **(L)** Mean ± standard deviation of frequency of events across all neurons between light (grey) and dark (black) conditions. **(M)** Mean ± standard deviation of fraction of active cells between light (grey) and dark (black) conditions. **(N)** Average calcium event across all neurons between light (grey) and dark (black) conditions. Shaded area represents the standard error of the mean. **(O)** Example head rotation tuning for an excitatory motor unit.

**Supplementary Fig. 3:**
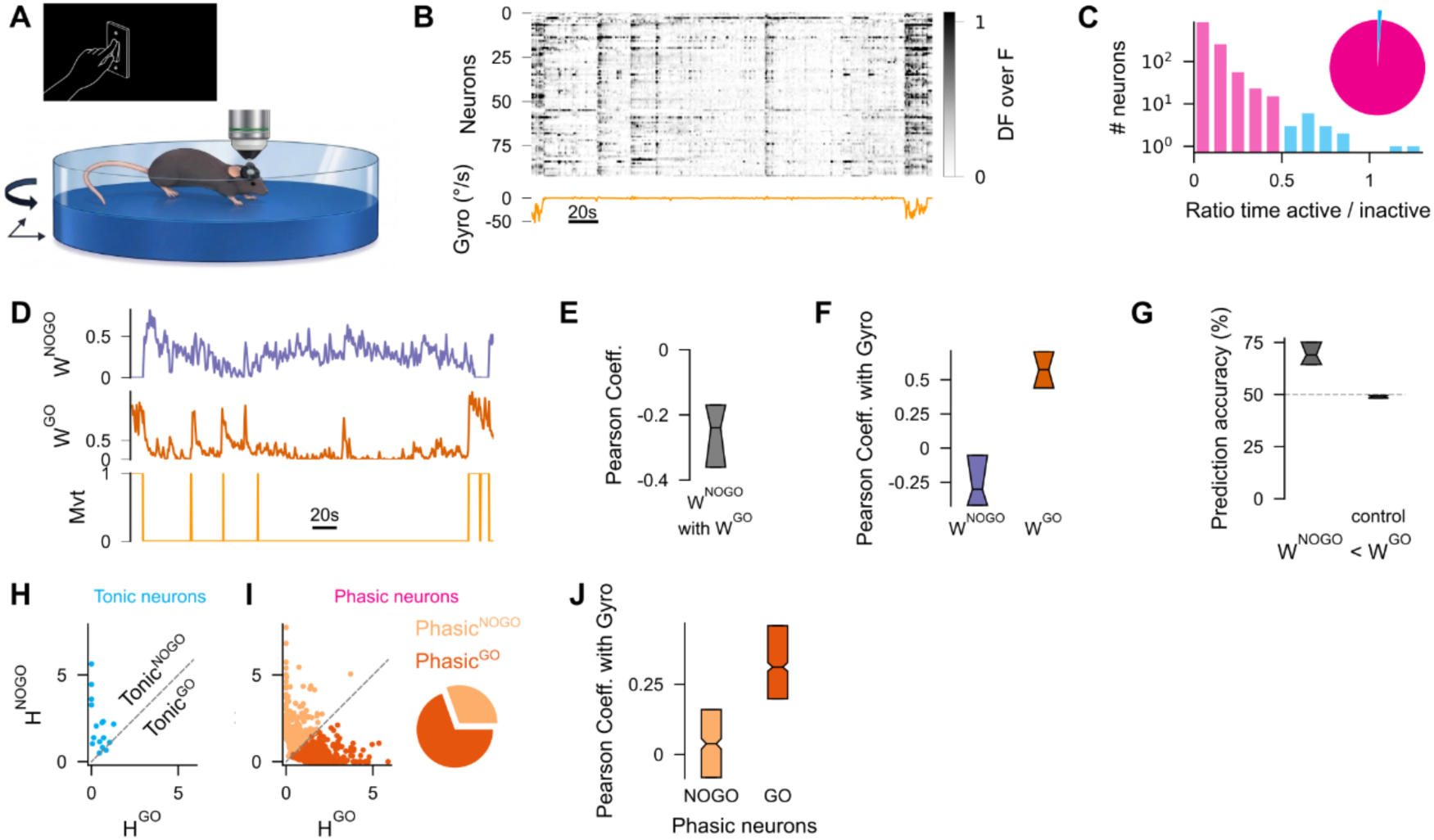
Two-photon in vivo imaging and NMF identify movement-related inhibitory activity profiles. **(A)** Experimental set up for *in vivo* two-photon calcium imaging of collicular neurons in response to orienting movements. Recordings were done in complete darkness. **(B)** Top: raster plot showing the variations in calcium fluorescence (DF over F) of VGAT^ON^ neurons for an example recording. Bottom: Synchronised recording of the floating platform rotation speed. **(C)** Histograms of the ratios between the number of DF over F timepoints above and below the midrange value. Cyan and magenta correspond to tonic and phasic neurons, respectively. Inset pie charts represent the percentage of tonic and phasic neurons in our recordings. Tonic neurons represented 1% ± 0.02 (mean ± standard deviation across recordings) of the total inhibitory population. **(D)** Example of the two temporal components, W^GO^ in purple and W^NOGO^ in orange, extracted by NMF from the recording in B. The bottom trace corresponds to the thresholded rotation speed of the platform to highlight periods of movement versus no movement. **(E)** Person correlation coefficients between the W^GO^ and W^NOGO^ components for all our recordings. Median [interquartile range]): −0.24 [−0.36, −0.17]. **(F)** Person correlation coefficients between the platform rotation speed and the W^GO^ and W^NOGO^ components. Median [interquartile range] across recordings; W^NOGO^ : −0.30 [−0.42, −0.05], W^GO^ : 0.57 [0.44, 0.71]. **(G)** Prediction accuracy achieved on test time points by comparing the value of the W^GO^ and W^NOGO^ components. Median [interquartile range] of cross validated prediction scores across recordings; data: 68.97 [64.23, 75.19]; scrambled data: 49.08 [48.29, 49.49]). Dotted line indicates the chance level. **(H)** H^NOGO^ and H^GO^ values for all tonic neurons. Dotted line represents the identity line which divides GO and NOGO neurons. **(I)** Same as H for phasic neurons. Pie chart on the right describes the percentages of Phasic^GO^ and Phasic^NOGO^ neurons among all inhibitory phasic neurons. Mean percentage of Phasic^GO^ among all phasic neurons ± standard deviation across recordings; 69.53 ± 12.32. **(J)** Person correlation coefficients between the platform rotation speed and the Phasic^GO^ and Phasic^NOGO^ neurons. Median [interquartile range]); Phasic^NOGO^: 0.04 [−0.08, −0.16], Phasic^GO^: 0.31 [0.20, 0.46].

**Supplementary Fig. 4:**
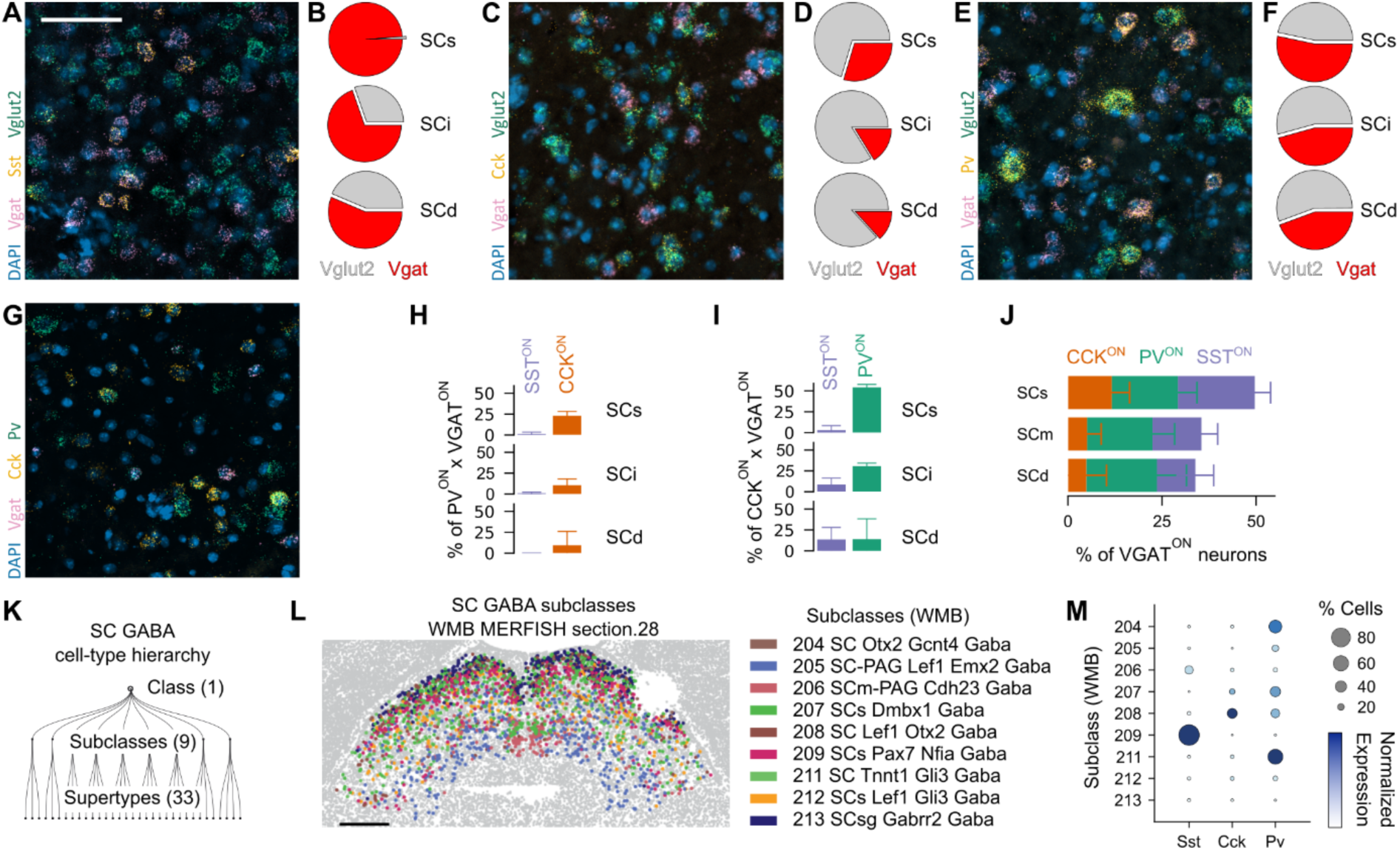
Sst, Cck and Pv expression in VGAT and VGLUT2 collicular neurons across layers. **(A)** Example field of view of a multiplexed FISH image of the SC when probing for Vgat, Vglut2 and Sst. **(B)** Pie charts describing the average percentages of SST positive neurons coexpressing Vgat (in red) and Vglut2 (in grey) within each SC layers. Mean ± standard deviation of percentage of Vgat positive cells among Sst positive cells across three coronal sections spanning the rostro-caudal axis of the SC; SCs: 99.2 ± 0.7; SCi: 69.6 ± 1.5; SCd: 56.0 ± 7.0. **(C-D)** Same as a-b probing for Cck instead of SST. SCs: 29.8 ± 3.2; SCi: 16.3 ± 3.4; SCd: 13.6 ± 6.1. **(E-F)**, Same as a-b, for Pv positive neurons. SCs: 53.1 ± 5.5; SCi: 45.7 ± 3.1; SCd: 44.4 ± 1.3. **(G)** Example field of view of a multiplexed FISH image of the SC when probing for Vgat, Cck and Pv. **(H)** Mean percentage of neurons coexpressing Sst or Cck together with Pv within the VGAT population for each SC layer. **i**, Mean percentage of neurons coexpressing Sst or Pv together with Cck within the VGAT population for each SC layer. **(J)** Mean percentages of Cck, Pv and Sst positive neurons among the VGAT positive population. Percentages values are stacked together and do not account for the overlap between subpopulations. Mean ± standard deviation. Pv; SCs: 17.6 ± 5.0, SCm: 17.3 ± 5.9, SCd: 18.7 ± 7.9. Sst; SCs: 20.4 ± 4.2, SCm: 13.0 ± 4.3, SCd: 10.2 ± 4.8. Cck; SCs: 11.7 ± 4.7, SCm: 5.1 ± 3.7, SCd: 4.9 ± 5.3. Scale bar represent 50 µm. Error bars represent the standard deviation. SCs: Superficial layers of SC; SCi: Intermediate layers of the SC; SCs: deep layers of the SC. **(K)** Allen Brain Cell (ABC) atlas taxonomy for SC GABA neurons (1 class) which can be subdivided in 9 subclasses and 33 supertypes. **(L)** Coronal section of the SC with all detected cells in gray and SC GABA neurons color-coded by their subclass belonging in the transcriptomic taxonomy. Note the dorsoventral layering of the different subclasses. Scale bar is 500 µm. **(M)** Dotplot illustrating mean scaled expression and fraction of expressing cells for Sst, Cck and Pv mRNA in different SC GABA subclasses as detected by scRNAseq. Note the specific enrichment of Sst expression in SC GABA subclass 209 and Cck expression in subclasses 207, 208 alongside Pv, which was instead more broadly expressed across different SC GABA subclasses.

**Supplementary Fig. 5:**
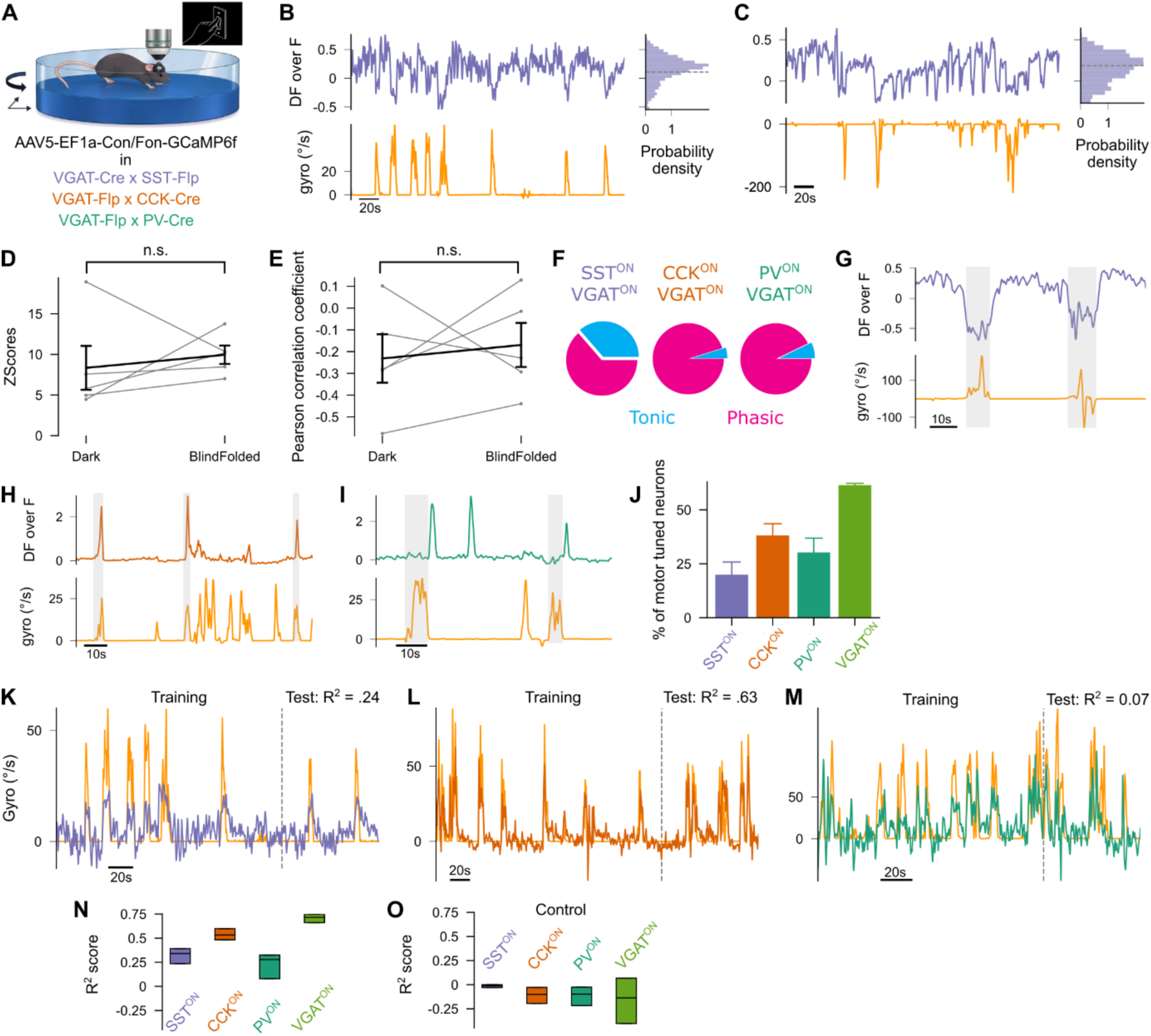
SSTON, CCKON and PVON neurons are differentially coupled to ongoing motor output. **(A)** Experimental set up for *in vivo* two-photon calcium imaging of SST^ON^, CCK^ON^ and PV^ON^ responses to orienting movements. **(B-C)** Example responses from the same SST^ON^ neuron recorded in the dark in A and in a blindfolded mouse in B. Top calcium traces. Bottom: synchronized platform rotation speed. Right: distribution of DF over F timepoints for the recording duration. **(D)** Paired values of ZScores for individual SST^ON^ neurons recorded in the dark versus in a blindfolded mouse. Mean ± standard deviation across neurons. Dark: 8.35 ± 5.40; Blindfolded: 9.96 ± 2.26. N = 5 neurons in 1 mouse. **(E)** Same as D for Pearson correlation coefficients. Dark: -.0.23 ± 0.22; Blindfolded: −0.17 ± 0.2. **(F)** Pie charts representing the percentage of tonic and phasic neurons for each inhibitory subpopulation. Mean of percentages of tonic neurons ± standard deviation across recordings; SST^ON^: 37.50 ± 48.41, CCK^ON^: 6.55 ± 9.44, PV^ON^: 13.26 ± 11.82. **(G-I)** Zoomed traces of the same example neurons shown in Fig. 2 B, C and D, respectively, to highlight the temporal relationship between each neuronal response and the movement of the mouse. **(J)** Percentage of motor tuned neurons among all recorded neurons for each inhibitory subpopulation. Error bars represent the standard deviation. Bootstrapped mean ± standard deviation: SST^ON^: 19.99 ± 11.70; CCK^ON^: 38.11 ± 10.91; PV^ON^: 30.25 ± 13.24; VGAT^ON^: 61.45 ± 1.73. **(K)** Example of prediction achieved of the platform rotation speed using SST^ON^ neurons. Please note that the curve in purple corresponds to a linear combination of the SST^ON^ neurons’ calcium traces. The calcium traces were tonically active and anticorrelated with the rotation speed and the linear model inverted the DF over F values to match the rotation speed of the platform. Dashed line represents how the data was divided into the training and test set. The R^2^ prediction score obtained from the test data is indicated above. **(L-M)** Same as K for CCK^ON^ and PV^ON^ neurons, respectively. **(N)** Boxplot of cross validated R^2^ prediction scores of the platform rotation speed obtained using the more predictive recording for SST^ON^, CCK^ON^ and PV^ON^ and VGAT^ON^ neurons. Median [interquartile range] across different train and test partitions randomly chosen over 1000 iterations. SST^ON^: 0.34 [0.24 0.39], CCK^ON^: 0.53 [0.48 0.60], PV^ON^: 0.28 [0.08 0.33], VGAT^ON^: 0.72 [0.66 0.75]. **o**, Same as o using instead circularly shift neuronal traces as a control. SST^ON^: −0.01 [−0.03 0.00], CCK^ON^: −0.10 [−0.19 −0.03], PV^ON^: −0.22 [−0.22 −0.02], VGAT^ON^: −0.14 [−0.40 0.07]. Boxplots show median and interquartile ranges.

**Supplementary Fig. 6:**
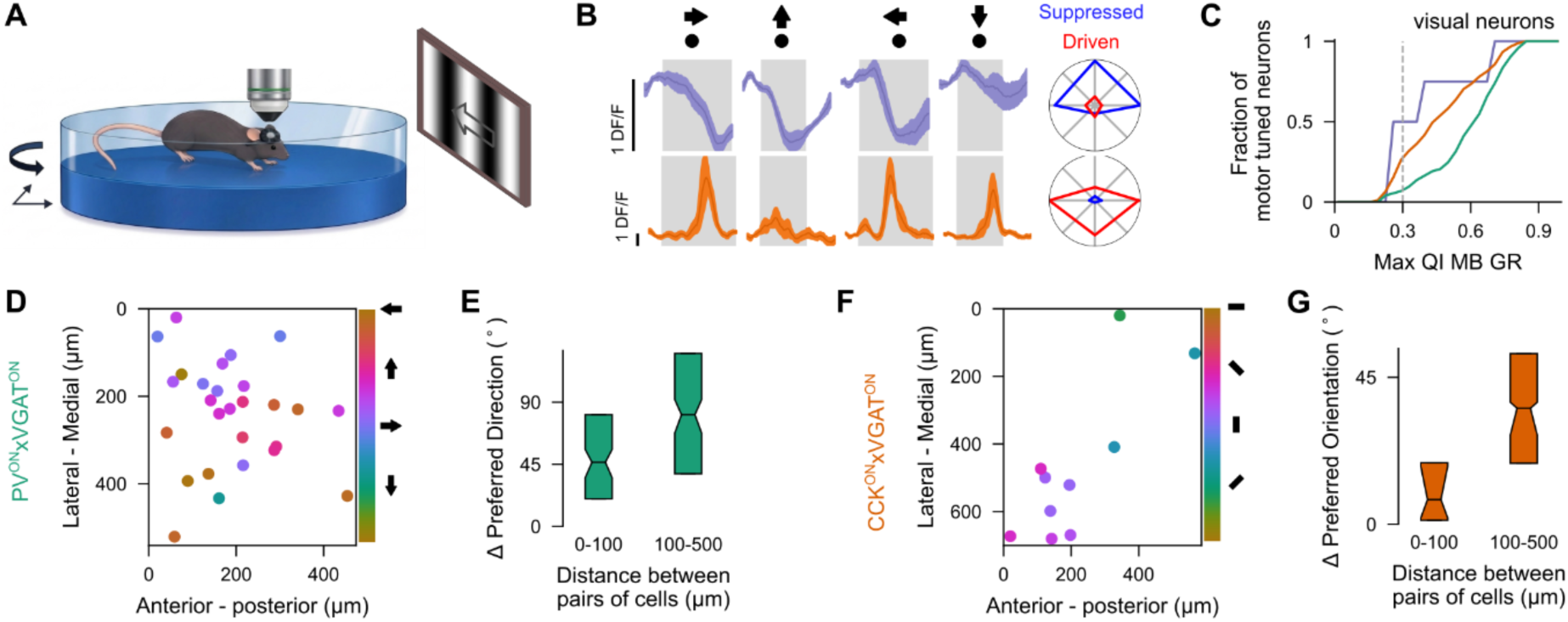
Visual tuning properties of SSTON, CCKON and PVON neurons. **(A)** Experimental set up for *in vivo* two-photon calcium imaging of SST^ON^, CCK^ON^ and PV^ON^ responses. **(B)** Example calcium responses to moving spots for SST^ON^ (top) and CCK^ON^ (bottom) neurons. Arrows above indicate the moving direction of gratings. Grey shade indicates the time the dots moved on the screen. Responses curves correspond to the mean responses across trials (dark line) surrounded by the standard error of the mean. Right polar histograms indicate the mean suppressed and driven responses for all moving directions. **(C)** Cumulative distributions of the maximum quality index in response to moving gratings and bars for each neuron within each inhibitory subpopulation. Dashed line indicates the threshold separating visual neurons from unresponsive neurons. One-tailed two sample Kolmogorov Smirnov test for QI^PV^ > QI^CCK^: Pvalue < 1.10^-^5. **(D)** Example recording field of view showing PV^ON^ DS neurons. Colour of dots represent the preferred direction of tuning according to the colour bar on the right. **(E)** Differences in preferred direction of tuning for pairs of PV^ON^ DS neurons located within a distance of 100µm from each other or further apart in the SC. Median [interquartile range] for pairs of DS neurons. Less than 100µm apart: 46.46 [19.96 80.93], N_neuron-pairs_ = 59. Further than 100 µm apart: 80.93 [38.17 125.43], N_neuron-pairs_ = 302. N_mice_ = 2, N_recordings_ = 4. **(F-G)** Same as D and E for the preferred orientations of CCK^ON^ OS neurons, respectively. Median [interquartile range] for pairs of OS neurons. Less than 100µm apart: 7.56 [1.22 18.85], N_neuron-pairs_ = 39. Further than 100 µm apart: 35.52 [18.68 52.28], N_neuron-pairs_ = 62. N_mice_ = 2, N_recordings_ = 8.

**Supplementary Fig. 7:**
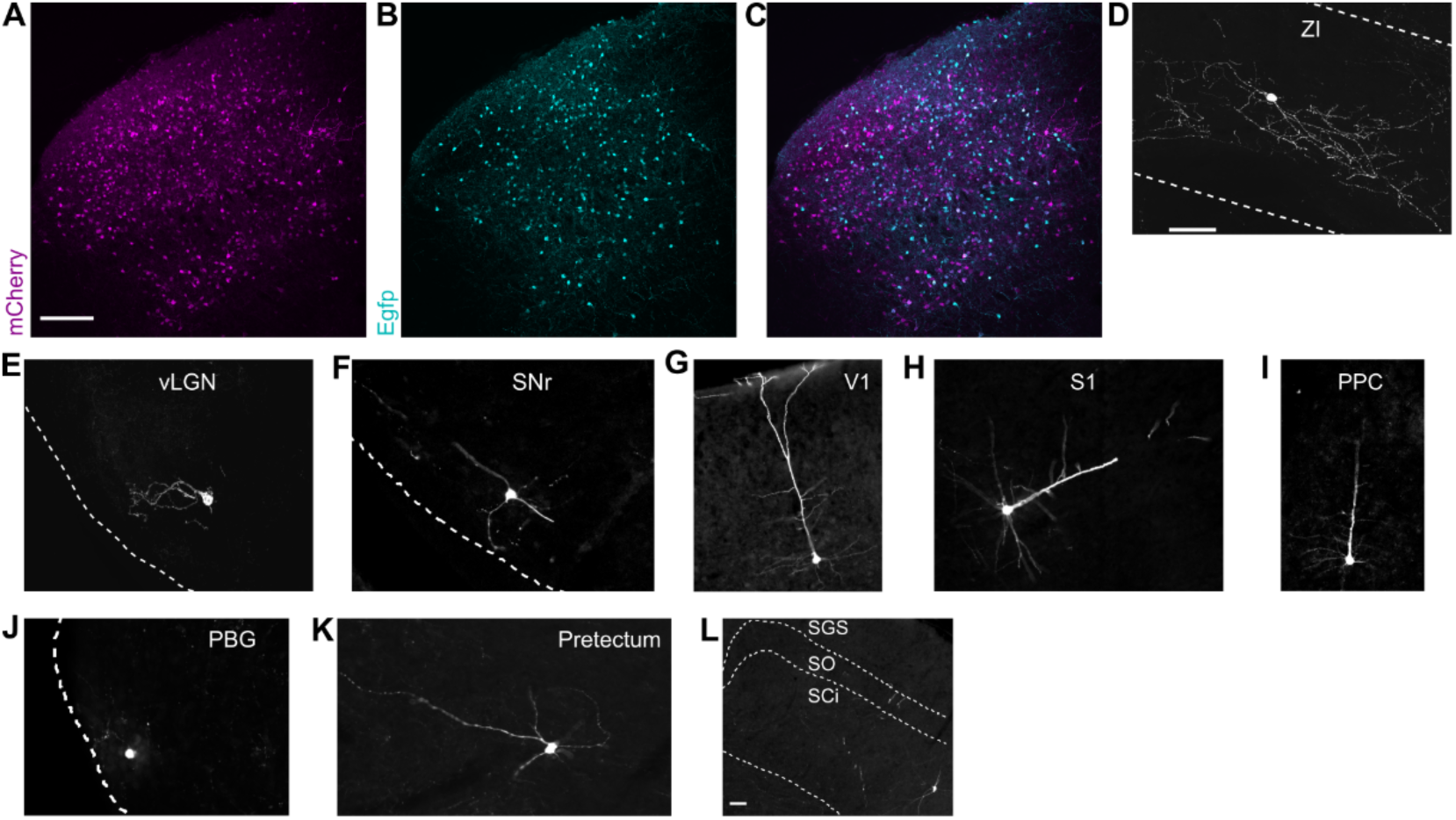
Brain-wide inputs to SSTON neurons. **(A-C)** Example field of view of the collicular injection site showing the expression of mCherry in A, Egfp in B and both of them combined in C. Scale bar is 200um. **(D-I)** Example of Egfp labelled cells found in the *Zona Incerta* (ZI, D), the ventral Lateral Geniculate Nucleus (vLGN, E), the *Substantia Nigra pars reticulata* (SNr, F), the Primary Visual cortex (V1, G), the Primary Somatosensory cortex (S1, H), the Posterior Parietal Cortex (PPC, I), the Parabigeminal nucleus (PBG, J), the pretectum (K) and contralateral intermediate layer of the SC (SCi, L). SGS: *Stratum Griseum Superficiale*. SO: *Stratum Opticum*.

**Supplementary Video 1: Prey capture in head-fixed conditions.**

**Supplementary Video 2: Moving-dot interception in freely moving conditions.**

